# Engineering CAR-Vδ2 T cells to boost persistence and anti-tumor function

**DOI:** 10.64898/2026.04.13.717612

**Authors:** Leonard Leong, Mansi Narula, Johannes Englisch, Cheryl Ou, Maksim Mamonkin, Norihiro Watanabe

## Abstract

Chimeric antigen receptor (CAR)-modified Vδ2 T cells are an attractive therapeutic cell platform for cancer immunotherapy. However, their clinical efficacy is limited by short *in vivo* persistence due to insufficient cytokine support and high susceptibility to activation-induced cell death (AICD). Through comparison of membrane-bound (mb) cytokines, we identified mbIL-18 to support superior anti-tumor activity of CAR-Vδ2 T cells *in vitro* and *in vivo*. To reduce constitutive surface exposure of IL-18 and enable antigen-driven signal 3, we fused MyD88 - the key IL-18R signaling mediator - to an extracellular domain of Fas (Fas88). Antigen stimulation-induced FasL engagement of Fas88 triggered IL-18 signaling while simultaneously protecting Vδ2 T cells from AICD. Fas88-armed human CAR-Vδ2 T cells produced superior yet stimulation-dependent *in vivo* expansion and functional persistence in xenograft models of hematologic and solid malignancies. Together, these findings highlight the importance of IL-18 signaling and AICD resistance for CAR-Vδ2 T cell activity, enabling a single-transgene modification to limit inflammatory risk and facilitate clinical translation.

## Introduction

Off-the-shelf cell therapies have many advantages over autologous therapies offering immediate treatment at a fraction of the cost. Current efforts are focused on improving the potency and clinical efficacy of such cell products. Vδ2 T cells, a major subset of γδT cells found in peripheral blood, exert cytotoxicity against cancer cells through the Vγ9Vδ2 TCR by recognizing conformational changes of BTN3A1/BTN2A1 molecules induced by phosphoantigens such as isopentyl pyrophosphate, accumulated in cancer cells due to dysregulated mevalonate pathway^1–3^. In addition, Vδ2 T cells express innate cytotoxic receptors such as NKG2D, DNAM-1, and FcγRIII (CD16), offering multi-pronged recognition and elimination of cancerous cells^4^. Similar to conventional αβT cells, Vδ2 T cells can be redirected with chimeric antigen receptors (CARs) to enhance tumor killing^5–7^. Moreover, Vδ2 T cells are naturally non-alloreactive as the invariant Vγ9Vδ2 TCR recognizes identical antigens in all humans thus minimizing the risk of graft-vs-host disease and making Vδ2 T cells highly attractive for off-the-shelf cell therapy.

Early clinical trials of Vδ2 T cell-based therapies demonstrated their safety but also illustrated shorter persistence of these cells compared to conventional αβT-cell based alternatives, limiting their clinical benefit^8,9^. The abbreviated lifespan of therapeutic Vδ2 T cells in patients could be attributed to two different mechanisms. First, Vδ2 T cells produced limited quantities of pro-survival cytokines, such as IL-2, and therefore were reliant on the presence of these factors in the environment^10^. Second, Vδ2 T cells are highly susceptible to activation-induced cell death (AICD)^11^ which restricts their persistence following TCR or CAR stimulation. These bottlenecks can be replicated in preclinical models where unmodified or CAR-engineered Vδ2 T cells produced meaningful anti-tumor activity only in high doses, with a low disease burden, or upon continuous supplementation with exogenous cytokines, such as recombinant IL-2^5,12–15^.

In this study, we engineer CAR-Vδ2 T cells to provide an ectopic cytokine signal, thereby creating a self-sustaining cell product. We identified IL-18 as a key cytokine supporting *in vivo* proliferation and the anti-tumor function of CAR-Vδ2 T cells. Further, we engineered a chimeric receptor that triggered IL-18 signaling upon antigen stimulation and protected CAR-Vδ2 T cells from AICD simultaneously, leading to sustained anti-tumor activity across diverse xenograft disease models. These findings illuminate key mechanisms limiting the clinical potency of CAR-Vδ2 T cells and offer actionable solutions that directly inform the next generation of clinical trials.

## Results

### Characterization of CAR-Vδ2 T cells expressing membrane-bound cytokines

Pro-survival cytokines IL-12, IL-15, and IL-18 promote Vδ2 T cell expansion and support anti-tumor function *in vitro*^16^, We first explored whether the ectopic expression of membrane-bound (mb) forms of these cytokines on Vδ2 T cells can overcome short in vivo persistence of Vδ2 T cells. We opted for membrane-tethered cytokines to contain their signaling to engineered Vδ2 T cells and their immediate environment and prevent over-production of soluble cytokines and potential systemic toxicity^17^(Fig.1a). We generated bicistronic γ-retroviral vectors encoding mb-cytokines and a second-generation CD19.CAR (19BBz) separated by a furin cleavage site and a T2A self-cleaving peptide (Fig.1b). The 4-1BB endodomain was chosen over CD28 or CD27 as it supported superior *in vivo* anti-tumor function of Vδ2 T cells^18^. Transgenic Vδ2 T cells were generated using an optimized manufacturing method (**Supplementary Fig.1a**) which we have previously demonstrated to enhance the *in vivo* function of CAR-Vδ2 T cells by reducing cellular senescence and apoptosis^18^. We detected robust surface expression of mbIL-12 and mbIL-18 and a somewhat reduced detection of mbIL-15 three days after transduction (Fig.1c). We also confirmed the signaling activity of mb-cytokines in Vδ2 T cells using flow cytometry. As shown in Fig.1d, expression of mbIL-12, mbIL-15, and mbIL-18 induced steady-state phosphorylation of STAT4, STAT5, and P38 MAPK, respectively. Of note, phosphorylation of STAT4 and p38 MAPK was detected in both transduced and non-transduced Vδ2 T cells indicating that mbIL-12 and mbIL-18 stimulated T cells both *in cis* and *in trans*. In contrast, phosphorylation of STAT5 was mainly confined to mbIL-15/19BBz-transduced T cells in culture, suggesting that mbIL-15 only signals *in cis*. Further, we detected a gradual downregulation of the CAR expression in mbIL-15/19BBz Vδ2 T cells over time whereas the CAR expression in other conditions was stable throughout the manufacturing process (Fig.1e and **Supplementary Fig.1b**). Further, the expression of mbIL-12 and mbIL-15 hindered *ex vivo* expansion of transgenic Vδ2 T cells in some donors (Fig.1f). We observed a negative correlation between mbIL-15/19BBz expression in Vδ2 T cells and their expansion (**Supplementary Fig.1c**), suggesting mbIL-15 inhibited Vδ2 T cell expansion in our system.

**Figure 1.**
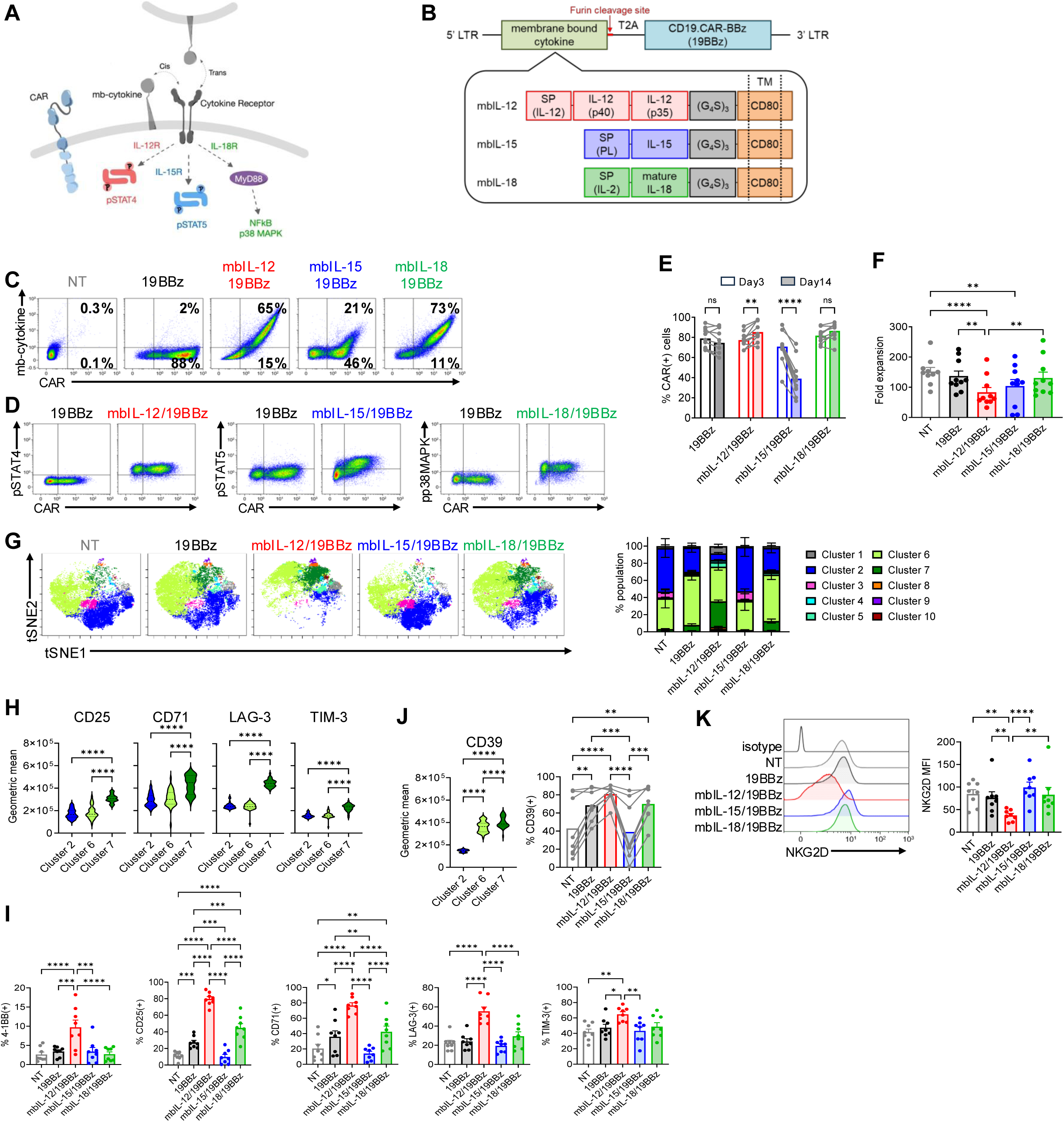
Characteristics of membrane-bound (mb) cytokines expressing CAR-Vδ2 T cells. **a,** Schematic of mb-cytokines and their downstream signaling. **b,** Schematic of γ-retroviral vectors encoding mb-cytokines and CD19.CARBBz. **c,** Representative flow plots of surface expression of CAR and mb-cytokines on day 3 post transduction from 11 independent donors. **d,** Representative flow plots showing phospho-flow of T cells from 3 independent donors. **e,** Surface CAR expression on day 3 and day 14 post transduction (mean ± S.E. n=11). **f,** Total cell fold expansion 14 days post transduction (mean ± S.E. n=9). **g,** Visualization of t-distributed stochastic neighbor embedding plots (viSNE) of concatenated events from 5 different cell products with 18 surface phenotype markers followed by FlowSOM metaclustering (cluster 1-10). viSNE plots (left) and bar graph summarizing % cluster in each product (right) (mean ± S.E. n=8). **h,** Geometric means of the indicated surface markers in cluster 2, 6 and 7 (mean ± S.E. n=8). **i,** Percent expression of the indicated surface markers on each Vδ2 T cell product (mean ± S.E. n=8). **j,** Geometric mean of CD39 expression in cluster 2, 6 and 7 (left) and percent expression of CD39 in each Vδ2 T cell product (mean ± S.E. n=8). **k,** A representative overlay histogram showing NKG2D expression from 8 independent donors (left) and bar graph summarizing the relative MFI in each product with multiple donors (mean ± S.E. n=8). Statistical significance is calculated by a two-way ANOVA with Sidak’s multiple comparison (e) or a one-way ANOVA with Tukey’s multiple comparison (f, h, i, j, k). ^✱^p ≤ 0.05, ^✱✱^p ≤ 0.01, ^✱✱✱^p ≤ 0.001, ^✱✱✱✱^p ≤ 0.0001, ns; non-significant.

Next, we evaluated whether expression of mb-cytokines affected the cell surface phenotype of CAR-Vδ2 T cells by assessing expression of activation markers, inhibitory receptors, senescence markers and memory phenotype simultaneously using multicolor spectral flow cytometry. Dimensionality reduction by the visualization of t-distributed stochastic neighbor embedding (viSNE) followed by a clustering by FlowSOM generated 10 metaclusters in concatenated data (**Supplementary Fig.1d**). As shown in Fig.1g, clusters 2 and 6 were dominant in NT, 19BBz, mbIL-15/19BBz and mbIL-18/19BBz Vδ2 T cells, with a largely similar cluster composition in NT and mbIL-15/19BBz, and 19BBz and mbIL-18/19BBz. In contrast, mbIL-12/19BBz showed an enrichment of cluster 7 compared to others. Cells in cluster 7 had a high expression of activation makers and inhibitory receptors CD25, CD71, LAG-3, and TIM-3 (Fig.1h and **Supplementary Fig.1d,e**) suggesting higher basal activation of mbIL-12/19BBz Vδ2 T cells during manufacturing (Fig.1i). CD39 expression differentiated clusters 2 and 6, with a higher expression of CD39 in 19BBz and mbIL-18/19BBz compared to NT and mbIL-15/19BBz, and the highest (albeit donor-dependent) expression in mbIL-12/19BBz Vδ2 T cells (Fig.1j). While we observed no differences in the differentiation/memory phenotype based on CD27 and CD45RA expression across different Vδ2 T cell products (**Supplementary Fig.1f**), NKG2D expression level was significantly lower in mbIL-12/19BBz Vδ2 T cells (Fig.1k), mirroring similar observation in NK cells^19^. Overall, we observed phenotypical similarity between NT and mbIL-15/19BBz, and 19BBz and mbIL-18/19BBz Vδ2 T cells, and a distinct phenotype for mbIL-12/19BBz Vδ2 T cells. Because mbIL-15/19BBz Vδ2 T cells invariably lost CAR expression at the end of manufacture, we excluded this construct from further comparisons.

### mbIL-18 supports CAR-Vδ2 T cell proliferation and enhances anti-tumor activity *in vitro*

We next evaluated whether the expression of mb-cytokine changed the cytokine prolife of CAR-Vδ2 T cells by stimulating them with NALM6 at a 1:1 effector-to-target ratio and measuring cytokines both intracellularly (following a 4-hr stimulation) and in the culture supernatant (following a 24-hr stimulation). mbIL-12/19BBz Vδ2 T cells had significantly higher intracellular production of IFNγ, TNFα, and GM-CSF in the absence of antigen stimulation (Fig.2a and **Supplementary Fig.2a**). Upon stimulation, all CAR-modified Vδ2 T cells produced IFNγ, TNFα, GM-CSF, and low levels of IL-2 (Fig.2a and **Supplementary Fig.2b**). Most CAR-Vδ2 T cells were IFNγ(+)TNFα(+) or IFNγ(+)TNFα(+)GM-CSF(+), demonstrating their ability to produce multiple cytokines (Fig.2a). Mirroring the intracellular staining, mbIL-12/19BBz Vδ2 T cells secreted high quantities of IFNγ, TNFα, soluble FasL (sFasL), and Granzyme B in the absence of antigen stimulation (Fig.2b). After a 24-hour stimulation, CAR-Vδ2 T cells armed with mbIL-12 or mbIL-18 secreted more IFNγ compared to unarmed CAR-Vδ2 T cells (Fig.2c). Those results indicate the expression of mbIL-12 increases the baseline activation of Vδ2 T cells in the absence of antigen resulting in elevated steady-state production of effector cytokines. By contrast, expression of mbIL-18 enhanced IFNγ production only upon antigen stimulation. After a 9-day coculture with NALM6 in the absence of exogenous cytokines (**Supplementary Fig.3**), mbIL-18/19BBz Vδ2 T cells showed superior functionality with robust CAR-Vδ2 T cell expansion and sustained suppression of tumor growth, while unarmed 19BBz Vδ2 T cells and mbIL-12/19BBz Vδ2 T cells had only transient anti-tumor activity and limited T cell expansion (Fig.2d-f). These data demonstrated the benefit of IL-18 over IL-12 signaling for sustained Vδ2 T cell expansion and cytotoxicity.

**Figure 2.**
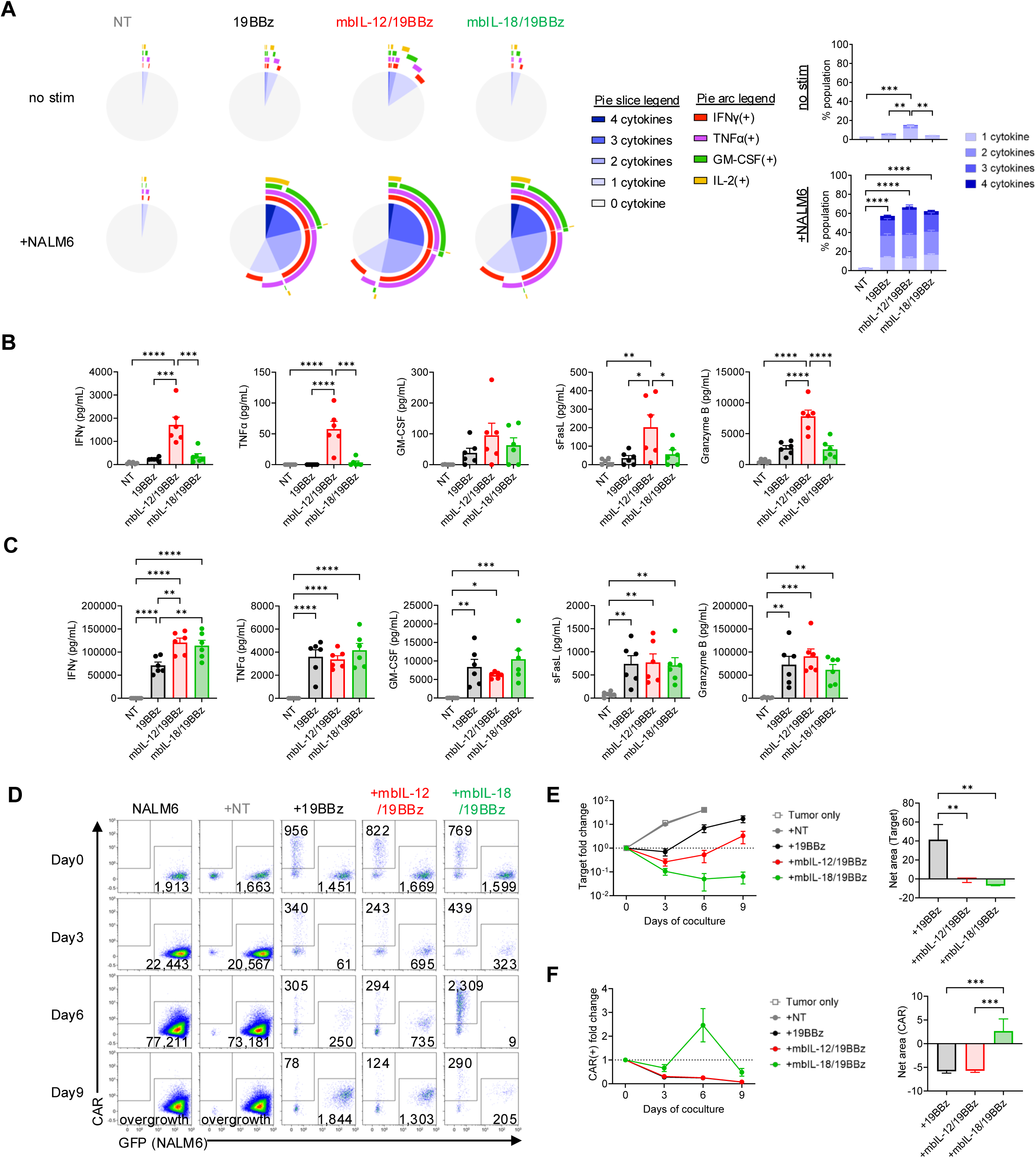
Effect of mbIL-12 and mbIL-18 on *in vitro* functionality of 19BBz Vδ2 T cells. **a,** Intracellular cytokine staining of each Vδ2 T cell product with or without antigen stimulation with NALM6 for 4 hrs. SPICE plots showing the expression of individual cytokines (left) and bar graph summarizing the frequency of cells secreting different number of cytokines in multiple donors (mean ± S.E. n=5). **b, c**, Cytokine secretion from each Vδ2 T cell product in the supernatant without stimulation **(b)** and with stimulation **(c)** for 24 hrs (mean ± S.E. n=6). **d-f,** Results of 9-day coculture experiments against NALM6. Representative flow plots of coculture experiments from 10 independent experiments **(d)**. Line graphs showing target cell fold change **(e)** and CAR(+) cell fold change **(f),** and bar graphs showing Net area (baseline = 1) from day 0 to day 9 for each mean ± S.E. n=10). Statistical significance is calculated by a one-way ANOVA with Tukey’s multiple comparison (a, b, c, e, f). ^✱^p ≤ 0.05, ^✱✱^p ≤ 0.01, ^✱✱✱^p ≤ 0.001, ^✱✱✱✱^p ≤ 0.0001.

### mbIL-18 augments *in vivo* proliferation and anti-tumor function of CAR-Vδ2 T cells in the absence of exogenous cytokines

We evaluated whether arming Vδ2 T cells with mb-cytokines would improve their *in vivo* functionality in an established NALM6 xenograft mouse model. We utilized a dual-luciferase imaging system in which NALM6 cells were modified to express a Click Beetle Green luciferase (CBG) while effector T cells were engineered to express an Aka Luciferase (Aka) thus enabling independent and simultaneous tracking of both cancer cells and effector T cells by IVIS imaging. In this experiment, mice received a single injection of freshly thawed Aka(+) Vδ2 T cells without exogenous cytokines (Fig.3a). The mbIL-12/19BBz Vδ2 T cells produced superior anti-tumor responses compared to the 19BBz Vδ2 T cell treatment, which showed only transient effects on time to recurrence (Fig.3b). Strikingly, administration of mbIL-18/19BBz Vδ2 T cells resulted in most potent *in vivo* anti-tumor effect long-term (Fig.3b) despite a slightly delayed response at early time points (**Supplementary Fig.4**), and significantly extended mouse survival (Fig.3c). This enhancement reflected superior expansion and persistence of mbIL-18/19BBz Vδ2 T cells compared to 19BBz and even mbIL-12/19BBz Vδ2 T cells which showed only limited expansion (Fig.3d,e). These results demonstrated that mbIL-18 arming augmented anti-tumor efficacy of CAR-Vδ2 T cells *in vivo* by enhancing their expansion and persistence in animals.

**Figure 3.**
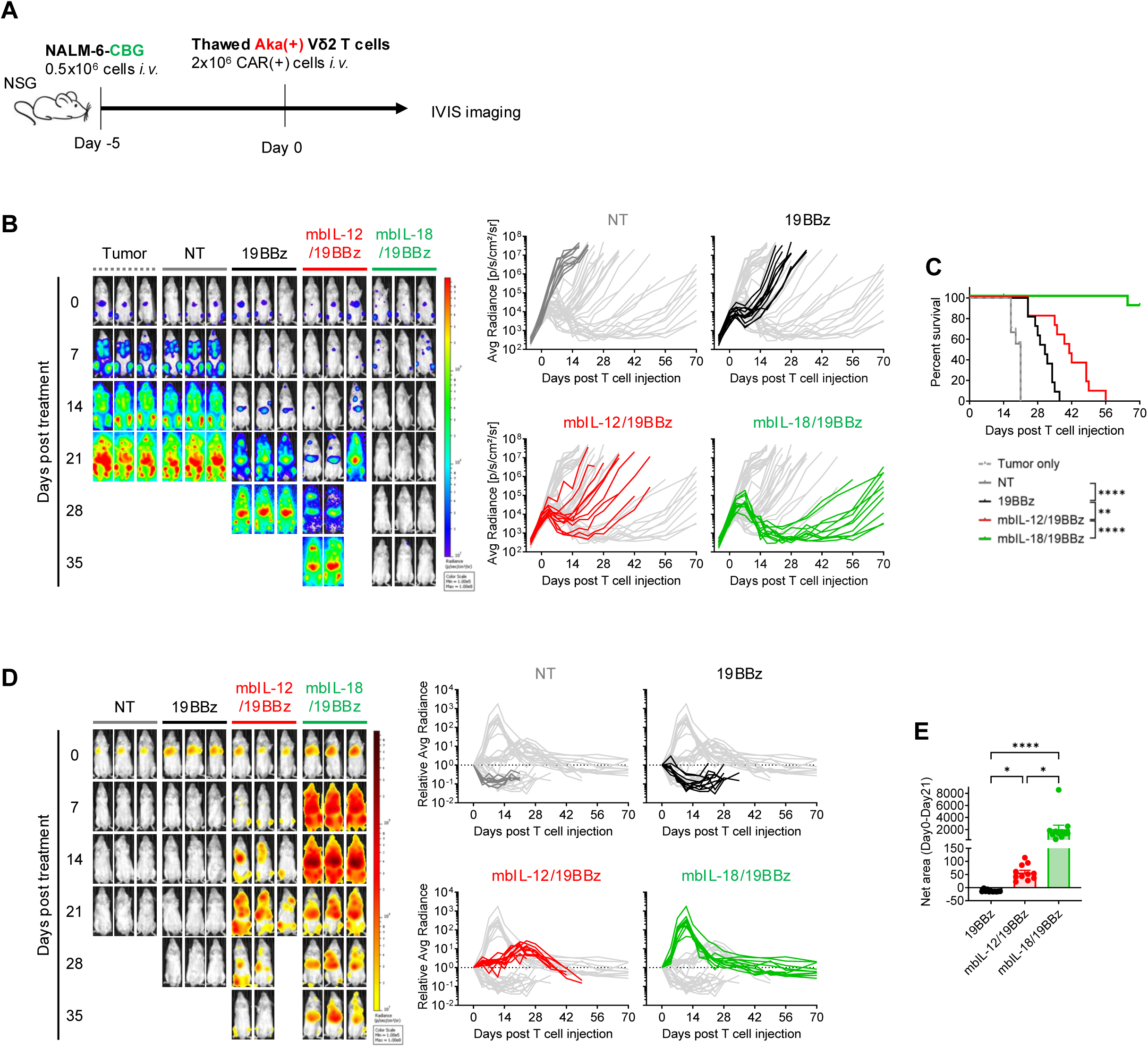
*In vivo* activity of mbCyto-19BBz Vδ2 T cells in a xenograft mouse model of human B-cell leukemia. **a,** Outline of *in vivo* experiments (n=7 in tumor only, n=9 in NT, n=11 in 19BBz and mbIL-12/19BBz, n=12 in mbIL-18/19BBz). **b,** Tumor progression detected by CBG bioluminescence. Mouse images show tumor bioluminescence at the indicated time points post Vδ2 T cell treatment and line graphs showing tumor progression in individual mice over time. **c,** Overall animal survival in each experimental group. **d,** Vδ2 T cell expansion detected by AkaLuc bioluminescence. Mouse images showing Vδ2 T cell levels at the indicated time points and line graphs showing Vδ2 T cell expansion and persistence in individual animals over time. **e,** Net area (baseline = 1) of AkaLuc bioluminescence fold change from day 0 to day 21. Statistical significance calculated by the log-rank test (c) or One-way ANOVA with Tukey’s multiple comparison (e). ^✱^p ≤ 0.05, ^✱✱^p ≤ 0.01, ^✱✱✱✱^p ≤ 0.0001.

### Expression of a truncated Fas receptor minimizes Fas-driven apoptosis in CAR-Vδ2 T cells but does not enhance long-term functional persistence

High sensitivity to activation-induced cell death (AICD), which is induced by Fas-FasL signaling, is another hallmark of *ex vivo* expanded Vδ2 T cells and limits their anti-tumor function^11^. Indeed, *ex vivo* expanded Vδ2 T cells showed high expression of Fas (Fig.4a) and expressed both membrane-bound and soluble FasL upon antigen stimulation (Fig.4b **and** Fig.2c). Therefore, we next investigated whether CAR-Vδ2 T cell functionality can be boosted by reducing their susceptibility to AICD. For that, we created a truncated Fas (ΔFas) construct which functions as a dominant-negative receptor that binds FasL without transmitting apoptotic signaling (Fig.4c). We expressed ΔFas from a gammaretroviral vector harboring ΔNGFR as a surrogate maker (Fig.4d). After transduction, we confirmed higher surface expression of Fas in the transduced population (NGFR+) compared to untransduced cells (NGFR-) by flow cytometry (Fig.4e). As expected, exposure to sFasL induced apoptosis in unarmed Vδ2 T cells but not in those expressing ΔFas, demonstrating functionality of the decoy receptor (Fig.4f). We next evaluated whether expression of ΔFas conferred functional advantage to 19BBz Vδ2 T cells in a 9-day coculture with NALM6. ΔFas arming boosted the anti-tumor effect of 19BBz Vδ2 T cells by enhancing their survival which resulted in an enrichment of the ΔFas-transduced population (Fig.4g). However, this *in vitro* observation did not translate *in vivo* as unarmed and ΔFas-armed 19BBz Vδ2 T cells produced largely similar anti-tumor activity in the NALM6 xenograft mouse model (Fig.4h,i). Overexpression of ΔFas did not boost long-term proliferation of Vδ2 T cells (Fig.4j), limiting survival advantage in mice treated with 19BBz/ΔFas Vδ2 T cells (Fig.4k). This suggested that the sequestration of death signal only was inadequate and necessitated additional support such as cytokines as shown before, to reinforce the potency of CAR-Vδ2 T cells.

**Figure 4.**
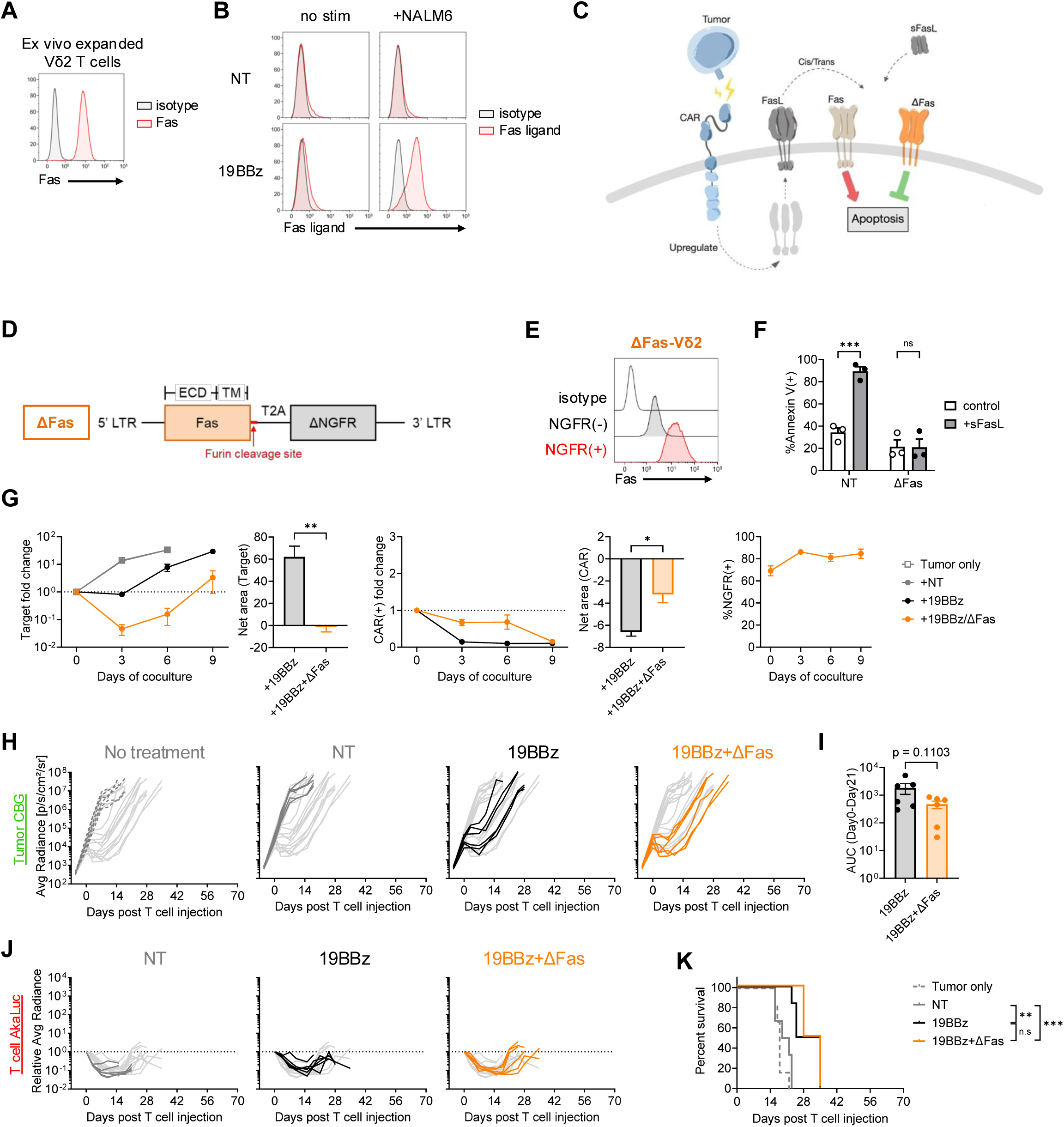
Overexpression of ΔFas on CAR-Vδ2 T cells improves short-term anti-tumor effect *in vitro* but not in an animal model. **a,** Representative histogram of Fas expression on *ex vivo* expanded Vδ2 T cells. **b,** Representative overlay histograms of Fas ligand expression on Vδ2 T cells with or without antigen stimulation for 24 hrs. **c,** Schematic of ΔFas expression and expected downstream events. **d,** Schematic of γ-retroviral vector encoding ΔFas. **e,** Representative overlay histogram of surface Fas expression in transduced (NGFR+) and untransduced (NGFR-) population in ΔFas-modified Vδ2 T cells. **f,** Bar graph showing % apoptotic cells (Annexin V+ cells) upon exposure to soluble Fas ligand for 24 hrs (mean ± S.E. n=3). **g,** Results of 9-day coculture experiments against NALM6. Line graphs showing target cell fold change (left), CAR(+) cell fold change (middle) and % NGFR(+) cells (right) over time, and bar graphs showing Net area (baseline = 1) of tumor fold change (left) and CAR(+) fold change (right) from day 0 to day 9 (mean ± S.E. n=5). **h-k,** Results from *in vivo* experiments in NALM6 xenograft mouse model (n=6/group). **h,** Tumor progression in individual mice over time detected by CBG bioluminescence. **i,** AUC of tumor bioluminescence from day 0 to day 21 post Vδ2 T cell treatment. **j,** Vδ2 T cell expansion in individual mice over time detected by AkaLuc bioluminescence. **k,** Overall animal survival in each experimental group. Statistical significance is calculated by a two-way ANOVA with Sidak’s multiple comparison (f), Paired t test (g), Unpaired t test (i), or the log-rank test (k). ^✱^p ≤ 0.05, ^✱✱^p ≤ 0.01, ^✱✱✱^p ≤ 0.001, ns; non-significant.

### A Fas-MyD88 fusion receptor induces IL-18 signaling in activated Vδ2 T cells while protecting from Fas-mediated apoptosis and boosting their anti-tumor function

While mbIL-18 significantly enhanced CAR-Vδ2 T cell function against the tumor *in vivo*, after an allogeneic OTS therapy with CAR-Vδ2 cells, trans-presentation of mbIL-18 to host immune cells can spur their activation and potentially accelerate rejection of the allogeneic product. Indeed, exposure to mbIL-18 enhanced the cytotoxicity of αβT cells expressing survivin-specific TCR^20^ when these were cocultured with BV173 cell lines expressing mbIL-18 (**Supplementary Fig.5a**). Therefore, we aimed to restrict the beneficial IL-18 signaling to therapeutic Vδ2 T cells without exposing the cytokine to the surrounding milieu. Further, we hypothesized that IL-18 signaling will synergize with AICD protection via Fas signaling diversion to additionally improve Vδ2 T cell function. Because IL-18R primarily signals through MyD88, we created a Fas-MyD88 fusion receptor (Fas88), to convert pro-apoptotic Fas signaling to pro-survival IL-18/MyD88 signaling, in response to antigen stimulation in CAR-Vδ2 T cells (Fig.5a,b). As expected, gammaretroviral expression of Fas88 increased surface expression of Fas in Vδ2 T cells (Fig.5c), and blocked FasL-induced apoptosis (Fig.5d). We measured MyD88 signaling from Fas88 in Jurkat reporter cells expressing GFP under an NFκB-sensitive promoter and engineered with either ΔFas or Fas88. As shown in Fig.5e, exposure to sFasL resulted in GFP expression only in Fas88-modified Jurkat reporter cells, demonstrating NFκB signaling downstream of the Fas88 fusion receptor. We also confirmed by Western Blot that Fas88 expression in Vδ2 T cells resulted in a phosphorylation of NFκB, IκK, and p38 MAPK - key mediators of IL-18 signaling - in the presence of soluble FasL (Fig.5f). In a 9-day coculture experiments, Fas88-armed 19BBz Vδ2 T cells produced superior cytotoxicity against NALM6 and mounted robust expansion compared to unarmed 19BBz Vδ2 T cells (Fig.5g). Fas88 had a similar effect in 19BBz Vδ2 T cells modified to express both mbIL-18 and ΔFas (Fig.5g). In the NALM6 xenograft mouse model, Fas88 arming endowed CD19.CAR-Vδ2 T cells with superior expansion and persistence, resulting in elimination of leukemia in mice, on a par with 19BBz Vδ2 T cells co-expressing mbIL-18 and ΔFas (Fig.5h,i).

**Figure 5.**
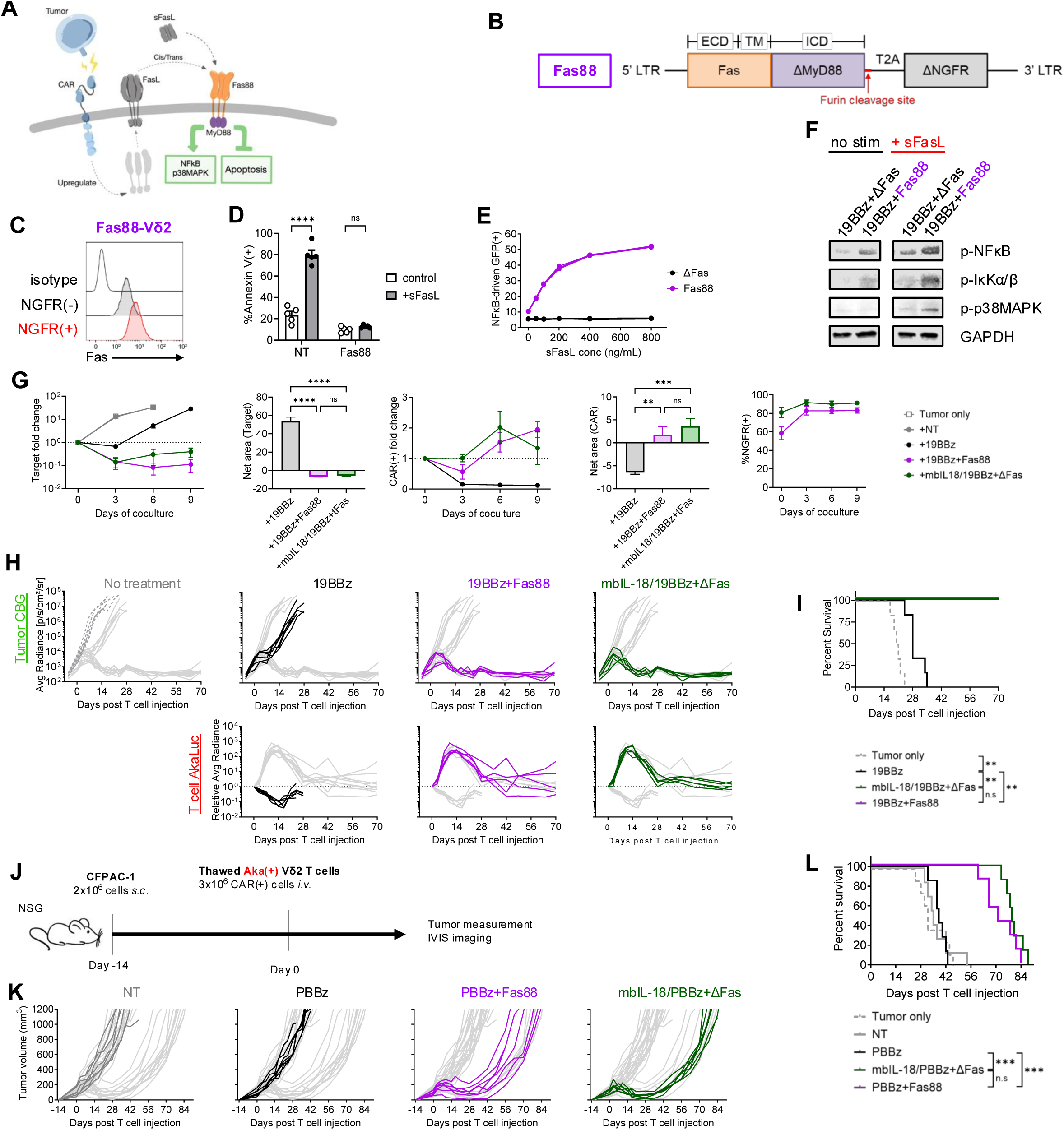
Fas88 confers AICD resistance and proliferative ability to CAR-Vδ2 T cells, enhancing their *in vivo* anti-tumor function. **a,** Schematic of Fas88 receptor and expected downstream events. **b,** Schematic of γ-retroviral vector encoding Fas88. **c,** Representative overlay histogram of surface Fas expression in transduced (NGFR+) and untransduced (NGFR-) population in Fas88-modified Vδ2 T cells. **d,** Bar graph showing % apoptotic cells (Annexin V+ cells) upon exposure to soluble Fas ligand for 24 hrs (mean ± S.E. n=5). **e,** Line graph showing GFP induction in Jurkat-NFκB reporter cell line with either ΔFas or Fas88 modification upon exposure to soluble Fas ligand for 24 hrs at indicated concentration. (n=3, technical replicate). **f,** Expression of indicated proteins with or without exposure to soluble Fas ligand for 1 hr by Western blot. **g,** Results of 9-day coculture experiments against NALM6. Line graphs showing target cell fold change (left), CAR(+) cell fold change (middle) and %NGFR(+) cells (right) over time, and bar graphs showing Net area (baseline = 1) of tumor fold change (left) and CAR(+) fold change (right) from day 0 to day 9 (mean ± S.E. n=5). **h,i,** Results from in vivo experiments in NALM6 xenograft mouse model (n=6 in no treatment and 19BBz, n=5 in 19BBz+Fas88 and mbIL-18/19BBz+ΔFas). **h,** Tumor progression (top) and Vδ2 T cell expansion (bottom) in individual mice over time detected by either CBG bioluminescence or AkaLuc bioluminescence, respectively. **i,** Overall animal survival in each experimental group. **j,** Schematic of CFPAC-1 *in vivo* experiments (n=8 in tumor only, n=7 in NT, PBBz, PBBz+Fas88, and mbIL-18/PBBz+ΔFas). **k,** Line graphs showing tumor volume in each treatment group over time. **l,** Overall mouse survival in each experimental group. Statistical differences are calculated by Two-way ANOVA with Sidak’s multiple comparison (d), One-way ANOVA with Tukey’s multiple comparison (g), or the log-rank test (i,l). ^✱✱^p ≤ 0.01, ^✱✱✱^p ≤ 0.001, ^✱✱✱✱^p ≤ 0.0001, ns; non-significant.

In addition to the systemic leukemia, we investigated the activity of Fas88-modified CAR-Vδ2 T cells in a model of a solid tumor. To that end, we generated Vδ2 T cells expressing a prostate stem cell antigen (PSCA)-specific CAR (PBBz) and evaluated their function in a pancreatic xenograft mouse model using subcutaneous human CFPAC-1 tumor (Fig.5j). In this model, unarmed PBBz Vδ2 T cells had no measurable anti-tumor activity, while Fas88 modification enhanced their anti-tumor effects with potency comparable to the mbIL-18 + ΔFas modification and extended animal survival although tumor relapses were eventually detected in all mice (Fig.5k,l). While the expansion of PBBz+Fas88 and mbIL-18/PBBz + ΔFas Vδ2 T cells in the liver at early time points (**Supplementary Fig.5b**), interfered with their detection at the subcutaneous tumor site due to the overlapping signal, the overall T cell signal was significantly higher than that of PBBz Vδ2 T cells which showed only minimal expansion (**Supplementary Fig.5c,d**). These results indicate that Fas88 modification conferred superior *in vivo* expansion to CAR-Vδ2 T cells enhancing their anti-tumor effect in both systemic leukemia and solid tumor models without exogenous cytokine support.

### Fas88 supports functional expansion of Vδ2 T cells expressing a CD28-costimulated CD5.CAR in a model of human T-ALL

Fas88 expression augmented the function of Vδ2 T cells expressing 4-1BB costimulated CARs so we sought a combinatorial benefit using CD28-costimulated CAR. We therefore co-expressed Fas88 with a 2nd generation CD5-targeting CAR with a CD28 costimulatory domain (5.28z) that has been clinically validated with autologous and donor-derived αβT cells targeting T cell malignancies^21,22^. Because CD5 is also expressed on the surface of Vδ2 T cells (**Supplementary Fig.6a**), modification of Vδ2 T cells with 5.28z caused excessive CAR signaling resulting in over-stimulation and/or fratricide and impairing cell expansion during manufacturing process (Fig.6a). To minimize unwanted CAR signaling during the *ex vivo* expansion, we utilized pharmacological tyrosine-kinase inhibitors (TKIs), dasatinib and ibrutinib, previously demonstrated to rescue the expansion of CD7.CAR-expressing αβT cells^23^ by blocking CD3ζ and CD28 signaling via Lck and Itk inhibition, respectively. As expected, TKI supplementation restored the expansion of 5.28z-expressing Vδ2 T cells comparable to control NT Vδ2 T cells (Fig.6a). Importantly, the expression of CAR and Fas88 were similar in T cells expanded with or without TKI addition, indicating retroviral transduction was not inhibited by TKIs (Fig.6b). We next investigated whether transgene expression and/or TKI supplementation affect the surface phenotypes of Vδ2 T cells by performing multicolor spectral flow cytometry followed by dimensionality reduction analysis and clustering (Fig.6c). The data split across 8 metaclusters demonstrated the enrichment of clusters 5-8 in Vδ2 T cells expanded without TKI (especially with 5.28z modification) whereas TKI supplementation resulted in an enrichment of cluster 1 cells regardless of the genetic modification (Fig.6c). Hierarchical clustering revealed that clusters 5-8 enriched in 5.28z Vδ2 T cells showed higher expression levels of activation markers (CD69, 4-1BB, CD25) and inhibitory receptors (CD39, TIGIT, TIM-3, LAG-3) compared to the condition with TKI (Fig.6d and **Supplementary Fig.6b**), likely reflecting antigen-driven tonic signaling through the CAR. In contrast, cluster 1 cells that were highly enriched upon TKI supplementation demonstrated a resting phenotype with reduced expression of activation/effector markers and a higher frequency of a central memory-like population compared to cells expanded without TKI (Fig.6d and **Supplementary Fig.6c**). Due to the poor expansion of 5.28z Vδ2 T cells without TKI, we only investigated the function of 5.28z Vδ2 T cells manufactured with TKI.

**Figure 6.**
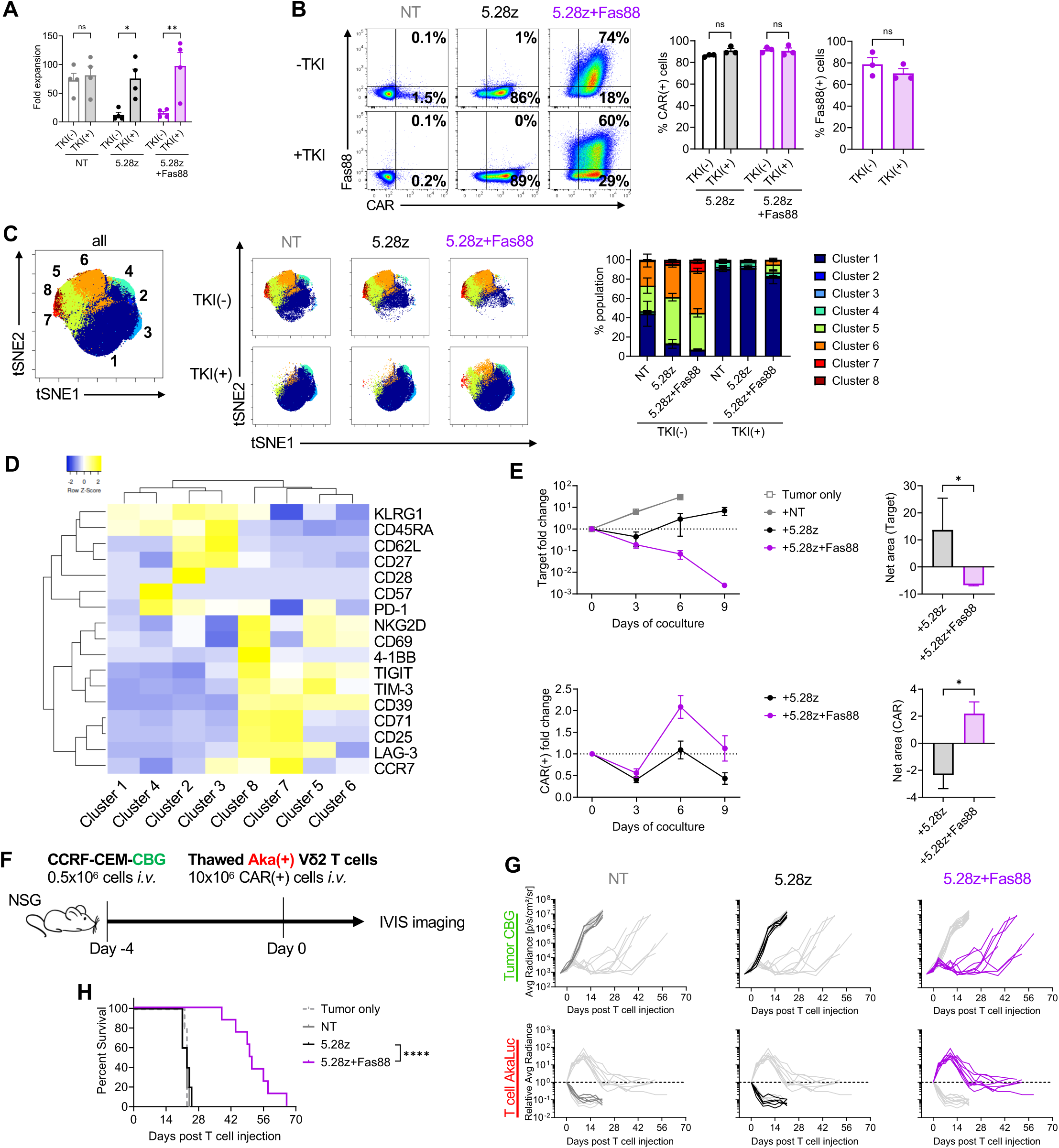
Generation and function of Fas88-armed CD5.CAR-Vδ2 T cells. **a,** Total expansion of cells generated with (+) or without (-) TKI on day 14 post-transduction (mean ± S.E. n=4). **b,** Representative flow plot (left) of surface expression of CAR and Fas88 (NGFR) in Vδ2 T cells +/− TKI on Day 14 post transduction and bar graphs summarizing multiple donor results (mean ± S.E. n=3). **c,** Visualization of t-distributed stochastic neighbor embedding plots (viSNE) of concatenated events from 6 different conditions (NT, CAR, CAR+Fas88, with or without TKI, 4 donors/condition) with 17 surface phenotype markers followed by FlowSOM metaclustering (cluster 1-8). Individual viSNE plots (left) and bar graph summarizing % cluster in each condition (right) (mean ± S.E. n=4). **d,** z-score heatmap of median expression of surface phenotype markers within 8 FlowSOM metaclusters with hierarchical clustering. **e,** Results of 9-day coculture experiments against CCRF-CEM (mean ± S.E. n=7). Line graph showing target cell fold change (top) and CAR(+) cell fold change (bottom) over time, and bar graphs showing Net area (baseline = 1) of tumor fold change (top) and CAR(+) fold change (bottom) from day 0 to day 9. **f,** Schematic of the T-ALL xenograft mouse model (n=5 in NT, n=5 in 5.28z, n=8 in 5.28z/Fas88). **g,** Tumor progression (top) and Vδ2 T cell expansion (bottom) in individual mice over time detected by either CBG bioluminescence or AkaLuc bioluminescence, respectively. **h,** Overall mice survival in each experimental group. Statistical differences are calculated by Two-way ANOVA with Sidak’s multiple comparison (a,b), Paired t-test (b), Wilcoxon-signed rank test (e), or log-rank test (h). ^✱^p ≤ 0.05, ^✱✱^p ≤ 0.01, ^✱✱✱✱^p ≤ 0.0001, ns; non-significant.

Upon a 9-day coculture with a CD5+ T cell acute lymphoblastic leukemia (T-ALL) cell line CCRF-CEM, unarmed 5.28z Vδ2 T cells transiently controlled tumor cell growth at day 3 but failed to sustain leukemia control (Fig.6e). In contrast, Fas88 armed 5.28z Vδ2 T cells expanded robustly and eliminated leukemia (Fig.6e). We then investigated the *in vivo* efficacy of 5.28z+Fas88 Vδ2 T cells in a CCRF-CEM T-ALL xenograft NSG mice model, using a dual-luciferase system to track tumor growth and T cell expansion simultaneously (Fig.6f). Unarmed 5.28z Vδ2 T cells injected i.v. produced no meaningful anti-tumor activity and did not expand *in vivo,* similar to control NT Vδ2 T cells. However, 5.28z+Fas88 Vδ2 T cells expanded in mice peaking on Day 7-10 post-infusion and mediated stronger tumor control compared to other groups (Fig.6g) resulting in a significant improvement in the overall animal survival (Fig.6h). Thus, Fas88 augmented functional persistence of CD5.CAR-Vδ2 T cells in an aggressive xenograft model of human T-ALL.

## Discussion

We describe a strategy to overcome the short lifespan of CAR-Vδ2 T cells – a major challenge limiting their *in vivo* activity^24^. Embedding IL-18 cytokine signaling within CAR-Vδ2 T cells increased their expansion and persistence resulting in an improvement in efficacy, underscoring the importance of cytokine support for a sustained activity post-infusion. Furthermore, coupling of the IL-18R signaling adaptor protein, MyD88, to a pro-survival Fas decoy provided an inducible cytokine signal to CAR-Vδ2 T cells combined with AICD protection for maximal anti-tumor effects. Together, these innovations establish a transformative framework for engineering highly durable and clinically potent CAR-Vδ2 T-cell therapies.

The potential of Vδ2 T cells (with or without CAR engineering) as tumor killers is well established^1^. Several studies have demonstrated the benefit of expanding or priming Vδ2 T cells with specific cytokines, alone or in combination, to boost cell proliferation and anti-tumor function^25,26^. Indeed, Vδ2 T cells are highly sensitive to polarization dependent on the cytokine milieu^27–29^. IL-12 and IL-18 promote Th1 Vδ2 T cell subsets^30–32^ while integrating IL-15 improved *ex vivo* proliferation and clinical efficacy^25^ of Vδ2 T cells. Built-in cytokine signals eliminated the need for continuous exogenous cytokine supplementation in “armored” CAR-αβT cells but only a few studies have used CAR-Vδ2 T cells. To our knowledge, a head-to-head comparison of CAR-Vδ2 T cells armed with different cytokines has not been published prior to this study.

In CAR-αβT cells, IL-12 has been shown to drive IFNγ production and promote anti-tumor activity^33–35^. We observed an upregulation of activation markers and effector cytokine secretion in mbIL-12-modified CAR-Vδ2 T cell product, resulting in a distinct phenotype compared to control CAR-Vδ2 T cells or those expressing mbIL-15 or mbIL-18. It is possible that a basal activated state in mbIL-12 CAR-Vδ2 T cells results in their hyperstimulation and premature cell death leading to a reduced expansion during manufacturing. Expression of soluble IL-15^7^ or an IL-15Rα/IL-15 fusion protein^36^ has been shown to enhance *in vivo* persistence and potentiate the anti-tumor function of Vδ2 T cells. In our system, we observed a significant loss of mbIL-15/CAR(+) cell fraction over the course of Vδ2 T cell generation, which has not been reported in previous studies. This could be due to the presence of human platelet lysate (HPL) in conditioning medium combined with constitutive mbIL-15 signaling. HPL has been reported to contain increased levels of TGF-β^18,37^, which can modulate the function of Vδ2 T cells, and interact with IL-15 resulting in a range of effects from an improvement of cytotoxicity^38,39^ to induction of a suppressive Vδ2 T cell subset^40^. However, TGF-β and IL-15 have not been previously evaluated in the context of CAR-Vδ2 T cells and therefore this angle requires additional investigation. Notably, the loss of CAR was only observed in mbIL-15- but not in mbIL-12- or mbIL-18-expressing CAR-Vδ2 T cells. IL-18 signaling in conventional CAR T cells has been rigorously tested in various preclinical models^41,42^, including immunocompetent models^43,44^, and IL-18 armed CAR T cells demonstrated high response rates against refractory tumors in patients (NCT04684563, NCT06017258)^45,46^. CAR-γδ T cells modified to secrete Granzyme B-cleavable IL-18 showed improved tumor control and extended mice survival in a xenograft model of breast cancer^47^. Similarly, we observed a functional upregulation of effector cytokines upon tumor stimulation and a greater *in vivo* mbIL-18/CAR-Vδ2 T cell expansion compared to mbIL-12/CAR-Vδ2 T cells in the B-cell leukemia xenograft mouse model. However, one of the potential drawbacks of expressing cytokines on the surface of allogenic cells is the risk of cytokine trans-presentation to bystander host immune cells in the vicinity, spurring additional inflammation^46^ and facilitating allo-rejection. This is especially important in the context of immune reconstitution post allogeneic cell therapy treatment. In fact, induction of IL-18 signaling and enhancement of cytotoxicity upon trans-presentation of mbIL-18 has been shown mechanistically in *ex vivo* expanded αβT cells^48^. In line with these reports, we observed i) IL-18 signaling in non-transduced T cells co-expanding with mbIL-18/19BBz cells, and ii) enhanced cytotoxicity of αβT cells against mbIL-18 expressing target cells.

High sensitivity to AICD is another facet of CAR-Vδ2 T cells that restrains their therapeutic potency^11^. Interfering with the Fas/FasL axis has been explored in αβT cells to prevent AICD and improve persistence. Truncated Fas-engineered TCR^49^ or CAR-αβT cells^50^, or Fas gene knock-out CAR-αβT cells indeed persist longer and produce better *in vivo* anti-tumor activity. Additionally, chimeric Fas-switch receptors employing costimulatory endodomains, such as those of the TNFR superfamily, have been extensively screened in αβT cells^51–53^, and proven effective in redirecting Fas signal to reinforce endodomain-dependent T cell function. In our study, while expression of ΔFas mitigated the Fas signal in CAR-Vδ2 T cells and transiently improved their activity, it failed to translate into a sustained benefit *in vivo*, rendering the need for other critical, complementary signals such as cytokine signaling.

As observed, IL-18 signaling provided a substantial expansion benefit to CAR-Vδ2 T cells and thus utilized to create a fusion receptor with Fas that leveraged the inducible nature of Fas-FasL interactions in T cells to provide conditional IL-18 signaling in CAR-Vδ2 T cells upon Fas ligation. Conditional, Fas-mediated activation of the IL-18 signaling can help prevent overstimulation of therapeutic T cells and synchronize the cytokine signal with Signals 1 and 2 triggered by a CAR. Confining the IL-18 signal to therapeutic T cells will help reduce immunogenicity and unwanted hyperinflammatory manifestations associated with paracrine IL-18 signaling. Further, the Fas88 construct is compact (~1kb) and can be easily combined with an antigen receptor, a safety switch, or other transgenes in a single vector. However, Fas-mediated apoptosis is an important regulatory mechanism in T cells and its subversion via the Fas88 construct poses a potential safety concern. We have not observed uncontrolled lymphoproliferation in Fas88 armed T cells *in vitro* or *in vivo* as Vδ2 T cells demonstrated transient expansion upon antigen stimulation and gradual contraction in all animals. Still, this concern can be defused by including additional safety measures^54^ such as reversible (e.g. drug-sensitive degrons)^55^ or irreversible (e.g. inducible Caspase 9)^56^ systems for the clinical translation of Fas88/CAR-Vδ2 T cells.

The functional benefit of Fas88 was observed with both 4-1BB- and CD28-costimulated CARs, as demonstrated in three separate tumor xenograft models, illustrating the versatility of the Fas88 modification regardless of the tumor type or CAR design. A CD5-targeted CAR has been successfully tested clinically in αβT cells, where it demonstrated safety and efficacy^22^, and the current data follows a similar pattern at least preclinically in CAR-Vδ2 T cells. This offers a straightforward translational path to Fas88/CAR-Vδ2 T cells. In summary, these findings establish the feasibility of engineering Vδ2 T cells for substantially improved functional persistence in both hematopoietic and solid tumor models, positioning this platform for subsequent clinical translation.

## Materials and Methods

### Donors and cell lines

Peripheral blood mononuclear cells (PBMCs) were obtained from healthy volunteers after informed consent under protocols approved by the Baylor College of Medicine Institutional Review Board (H-45017). 293T (human embryonic kidney cell line), NALM6 (pre-B-ALL cell line), CFPAC-1 (pancreatic ductal adenocarcinoma), CCRF-CEM (T-ALL) were obtained from the American Type Culture Collection (Rockville, MD), and BV173 (B-cell precursor leukemia) from Leibniz Institute DSMZ-German Collection of Microorganisms and Cell Cultures GmbH (Braunschweig, Germany). 293T cells were maintained in DMEM (Dulbecco’s Modified Eagle Medium: Gibco BRL Life Technologies, Inc., Gaithersburg, MD) supplemented with 10% heat-inactivated fetal bovine serum (FBS) (Gibco BRL Life Technologies, Inc.) and 2 mM L-GlutaMAX (Gibco BRL Life Technologies, Inc.). NALM6, CCRF-CEM, and BV173 cells were maintained in RPMI-1640 (HyClone Laboratories, Marlborough, MA) supplemented with 10% FBS and 2 mM L-GlutaMAX. CFPAC-1 cells were maintained in IMDM (Iscove’s Modified Dulbecco’s Medium: Gibco) supplemented with 10%FBS and 2 mM L-GlutaMAX. Cells were maintained in a humidified atmosphere containing 5% carbon dioxide (CO_2_) at 37 °C and were routinely tested for Mycoplasma.

### Generation of viral vectors and virus production

The γ-retroviral vectors (SFG) encoding second-generation CAR constructs targeting CD19 with 4-1BB/CD3ζ signaling domain (19BBz) and CD5 with CD28/CD3ζ signaling domain (5.28z) were previously generated^18,57^. Second-generation CAR targeting PSCA with 4-1BB/CD3ζ signaling domain (PBBz) was cloned based on the CAR-PSCA.CD28z construct that was previously optimized^58^, by replacing CD28 costimulatory domain with 4-1BB by utilizing NEBuilder HiFi DNA Assembly (New England Biolabs, Ipswich, MA). To generate membrane-bound cytokine-encoding bicistronic retroviral vector with 19BBz, we first prepared gene fragments as follows: IL-12 – human IL-12p70 was PCR amplified from an expression vector (kindly provided by Dr. Masataka Suzuki at Baylor College of Medicine); IL-15 – human IL-15 (Uniprot: P40933) was gene-synthesized with preprolactin signal peptide^59^ (Integrated DNA Technologies, Coralville, IA); IL-18 – human IL-18 mature form (Uniprot: Q14116) was gene-synthesized with a human IL-2 signal peptide^43,44^; membrane anchor – a fragment comprising a (G_4_S)_3_ linker, CD80 transmembrane domain^60,61^ (amino acid position: 236-270, Uniprot: P33681), Furin cleavage site (RAKR)^62^, and T2A sequence was gene-synthesized. Each cytokine fragment and membrane anchor fragment were cloned into a retroviral vector encoding 19BBz. The mbIL-18 fragment was also cloned into a retroviral vector encoding F8 (FLAG epitope tag linked with G_4_S linker and CD8 stalk/transmembrane domain) generated in our laboratory. To generate truncated Fas and Fas88 construct, a Fas gene containing signal peptide, extracellular domain, and transmembrane domain was amplified from cDNA of activated T cells, and a truncated MyD88 sequence lacking C-terminal TIR domain^63^ was gene-synthesized. Either truncated Fas gene fragment only or truncated Fas and MyD88 were assembled into a retroviral vector encoding ΔNGFR. To generate a lentiviral vector encoding NFkB-GFP reporter, the sequence encoding NFkB response element to GFP was PCR amplified from pSIRV-NF-kB-eGFP (Addgene: #118093) and cloned into a lentiviral vector encoding Q8 (CD34 epitope tag linked with G_4_S linker and CD8 stalk/transmembrane domain) as a surrogate marker. The γ-retroviral vector encoding survivin-specific TCR^20^ (previously published; kindly gifted by Dr. Bilal Omer at Baylor College of Medicine), a click beetle green luciferase co-expressing green fluorescent protein (CBG/GFP), and Aka luciferase co-expressing GFP (Aka/GFP) (kindly gifted by Dr. Pradip Bajgain at NCI) were previously published^20,64^. We modified the Aka/GFP vector to replace GFP with ΔEGFR, which was PCR-amplified from the DNA template (kindly provided by Dr. David Quach at Baylor College of Medicine) to facilitate isolation of transduced population. Lentiviral supernatant and γ-retroviral supernatant were generated as previously described^65,66^.

### Generation of gene-modified Vδ2 T cells and cell lines

Gene-modified Vδ2 T cells were generated using our optimized manufacturing protocol as previously described^18^, with minor modifications. Briefly, PBMCs (10^6^ cells/mL) were stimulated with 1 μM of Zoledronate (Zoledronic acid; Sigma-Aldrich, St. Louis, MO) in the presence of 100 U/mL of recombinant human IL-2 (NIH, Bethesda, VA) in CTL medium composed of 47.5% RPMI-1640, 47.5% Clicks medium (FUJIFILM Biosciences, Santa Ana, CA) and 2 mM L-GlutaMAX supplemented with 5% of human platelet lysate (HPL; nLiven PR™, Sexton Biotechnologies, BioLife Solutions Inc., Bothell, WA). Five days later, γδT cells were negatively isolated using TCRγ/δ+ T Cell Isolation Kit (Miltenyi Biotec Inc., San Diego, CA) and cryopreserved. For retroviral gene transfer, isolated γδT cells were thawed and subjected to retroviral transduction with Retronectin reagent (Takara Bio USA, Inc., San Jose, CA) as previously described^18^. Cells were split and fed every 2–3 days with fresh CTL media plus IL-2 (100 U/mL). For CD5.CAR-Vδ2 T cells, Dasatinib (200 nM) (#S1021; Selleckchem, Houston, TX) and Ibrutinib (200 nM) (#S2680; Selleckchem) were added on the day of transduction and cells were split and fed every 2–3 days with fresh CTL media with IL-2, Dasatinib, and Ibrutinib. On day 14 post-transduction, cells were harvested and cryopreserved. For *in vivo* experiments, Aka/ΔEGFR(+) cells and CAR(+) cells were enriched, and αβTCR(+) cells and Vδ1TCR(+) cells were depleted using biotin-conjugated monoclonal antibody followed by anti-biotin microbeads (Miltenyi Biotec Inc.) using the MACS system (Miltenyi Biotec Inc.), during the manufacturing process. To generate cell lines overexpressing transgene, we used the same protocol as described above and isolated the transduced population by using either a cell sorter (SH800S, Sony Biotechnology, San Jose, CA) or MACS system. While gene-modified Vδ2 T cells were generated in CTL medium supplemented with 5% HPL, all *in vitro* functional assays were performed in CTL medium supplemented with 10% FBS.

### Flow cytometry

To investigate cell surface phenotype, cells were stained with fluorochrome-conjugated antibodies for 20 min at room temperature. All samples were acquired on a Gallios Flow Cytometer (Beckman Coulter Life Sciences, Indianapolis, IN) or Cytek Northern Lights (Cytek Biosciences, Inc., Fremont, CA), and data was analyzed using Kaluza 2.1 Flow Analysis Software (Beckman Coulter Life Sciences), FlowJo 10.9.0 (BD Biosciences), or Cytobank (Beckman Coulter Life Sciences). Heat map was generated using the Heatmapper software^67^. The Vδ2TCR(+) population in NT Vδ2 T cells and the CAR(+)Vδ2TCR(+) population in CAR-modified Vδ2 T cell products were analyzed for cell surface phenotype. Antibodies used in this study are listed in Supplementary Data 1.

### Phospho-flow cytometry

Gene-modified Vδ2 T cells were rested in 10% FBS-supplemented CTL medium without exogenous cytokine overnight prior to phospho-flow cytometry analysis. Cells were harvested and stained for surface antigens and BD Horizon Fixable Viability Stain 700 (FVS700) (#564997; BD Biosciences) at room temperature for 20 min. After washing, cells were fixed with 2% formaldehyde solution (F1635, Sigma-Aldrich) at room temperature for 10 min, washed, permeabilized with pre-chilled 100% methanol (Fisher Scientific, Pittsburgh, PA) for 30 min on ice, and then washed three times. Cells were then stained with the phospho-antibody for 60 min at room temperature. After incubation, cells were analyzed using a flow cytometer.

### Detection of surface Fas ligand expression

Gene-modified Vδ2 T cells were incubated with or without NALM6-CBG/GFP at a 1:1 ratio (0.2 x 10^6^ cells each) in the presence of Fas-blocking mAb (1 μg/mL) (#684401; BioLegend) and Batimastat (2 μM) (#SML0041; Sigma-Aldrich) for 24 hrs in 200 μL in a 96-well flat-bottom plate. After incubation, cells were harvested, stained and analyzed using a flow cytometer.

### Detection of apoptotic cells

Gene-modified Vδ2 T cells were incubated with or without soluble Fas ligand (800 ng/mL) (#589406; BioLegend) for 24 hrs. After incubation, cells were harvested, stained with Pacific Blue-conjugated Annexin V (#640918; BioLegend) and 7-AAD (#559925; BD Biosciences), then analyzed using a flow cytometer.

### Cytokine quantification

For intracellular cytokine staining, freshly thawed 0.2 x 10^6^ Vδ2 T cells were cocultured with 0.2 x 10^6^ NALM6-CBG/GFP in 200 μL in a 96-well flat-bottom plate in the presence of Monensin (BD GolgiStop, BD Biosciences, San Jose, CA) for 4 hrs. After the incubation, cells were stained for cell surface markers and FVS700. Stained cells were then fixed and permeabilized with BD Cytofix/Cytoperm (BD Biosciences) according to the manufacturer’s protocol. They were intracellularly stained for IFNγ, TNFα, GM-CSF, and IL-2 for 30 min at 4°C and analyzed using a flow cytometer. To measure cytokines secreted from gene-modified Vδ2 T cells, Vδ2 T cells were cocultured with NALM6-CBG/GFP as described above for 24 hrs. Supernatants were collected and stored at −80°C. Cytokine levels were measured by Human Luminex Discovery Assay (Merck Millipore, Billerica, MA) and samples were acquired by Luminex 200 instrument (Thermo Fisher Scientific, Invitrogen, Grand Island, NY) according to the manufacturer’s instructions.

### Coculture experiments

In the Vδ2 T cell coculture experiments with the tumor, freshly thawed 10,000 CAR(+) CAR-Vδ2 T cells were cocultured with 20,000 CBG/GFP(+) target cell lines in 200 μL without exogenous cytokine in a 96-well flat-bottom plate. In the coculture experiments with survivin-TCR-modified αβT cells, 2,500 survivin-TCR(+) T cells were cocultured with 20,000 BV173-CBG/GFP(+) cells with or without mbIL-18 modification in 200 μL without exogenous cytokine in a 96-well round-bottom plate. Half of the medium was replaced with fresh medium every 3 days. Cells were harvested, stained, and analyzed using a flow cytometer at the indicated time points. To quantify cell numbers by flow cytometry, 10 μL/sample of CountBright Absolute Counting Beads (Thermo Fisher Scientific) was added, and 7-AAD was added to exclude dead cells. 1,000 counting beads per sample were acquired.

### NFκB reporter assay

The gene-modified Jurkat reporter cell line was incubated with various concentrations of soluble Fas ligand for 24 hrs. After incubation, cells were harvested and analyzed for GFP induction using a flow cytometer.

### Western blots

Gene-modified Vδ2 T cells were stimulated with or without soluble Fas ligand (800 ng/mL) for 1 hour. After incubation, cells were harvested, and cell lysates were prepared using RIPA lysis buffer (#R0278; Sigma-Aldrich) and Halt™ Protease and Phosphatase Inhibitors (#78425; Thermo Scientific) for 30 mins on ice. Cell lysates were passed through a fine needle to shear genomic DNA. For gel electrophoresis, cell lysate samples were loaded onto Mini-PROTEAN® TGX™ Precast Gels (#4561036; Bio-Rad, Hercules, CA) under reducing conditions. Western blot was performed by wet transfer to nitrocellulose membranes. Membranes were incubated with the primary antibodies in 1% BSA/PBST overnight at 4°C. Then, they were incubated with corresponding luminescent secondary antibodies for 1 hr at room temperature before visualization on the LI-COR Odyssey imaging system (LI-COR Biotechnology, Lincoln, NE). Antibodies used for western blot are also listed in Supplementary Data 1.

### In vivo studies

Breeder pairs of NOD.Cg-Prkdc^scid^ Il2rg^tm1Wjl^/SzJ mice (NSG mice; Strain no. 005557) were purchased from the Jackson Laboratory and bred in the Baylor College of Medicine animal facility. Both female and male littermates (aged 8–12 weeks) were used for experiments. All animal experiments were conducted in compliance with the Baylor College of Medicine Institutional Animal Care and Use Committee (IACUC) (protocol no. AN-4758). To evaluate the in vivo anti-tumor effect of gene-modified Vδ2 T cells against NALM6, 0.5 x 10^6^ NALM6-CBG/GFP cells were injected into NSG mice intravenously. Five days later, freshly thawed 2 x 10^6^ AkaLuc(+) gene-modified Vδ2 T cells were injected intravenously. Both tumor cell growth and T cell expansion/persistence were evaluated by a dual-luciferase bioluminescence imaging system optimized in our laboratory. Briefly, mice were injected with 100 μL TokeOni (1 mM: Sigma-Aldrich) intraperitoneally and imaged for AkaLuc bioluminescence using an IVIS Lumina III imaging system (Caliper Life Sciences, Hopkinton, MA, USA). Subsequently, mice were injected with 100 μL D-luciferin (30 mg/mL; PerkinElmer, Waltham, MA, USA) and imaged with a 520 nm filter for CBG bioluminescence. For the subcutaneous tumor model, NSG mice were engrafted with 2 x 10^6^ CFPAC-1 subcutaneously and once tumors reached approximately 100 mm^3^ (day 14), freshly thawed 3 x 10^6^ AkaLuc(+) gene-modified Vδ2 T cells were injected intravenously. Tumor size was measured using calipers and tumor volume was calculated as follows: tumor volume (mm^3^) = length x width^2^ / 2. T cell expansion/persistence was evaluated by IVIS imaging with TokeOni injection as described above. For systemic CCRF-CEM model, 0.5 x 10^6^ CCRF-CEM-CBG/GFP cells were injected into NSG mice intravenously, and 4 days later, freshly thawed 10 x 10^6^ AkaLuc(+) gene-modified Vδ2 T cells were injected intravenously. Both tumor and T cells were tracked by dual-luciferase bioluminescence imaging as described above. Data was analyzed by Living Image software (Caliper Life Sciences).

### Statistical analysis

Statistical analysis was performed using GraphPad Prism 10 (GraphPad Software, La Jolla, CA, USA). The statistical tests applied in each experiment are described in the corresponding figure legends.

## Acknowledgements

We thank Dr. Malcolm Brenner and Dr. Helen Heslop at the Baylor College of Medicine for providing feedback on the manuscript, Dr. Bilal Omer at the Baylor College of Medicine for kindly gifting the γ-retroviral vector for Survivin TCR, Dr. Pradip Bajgain at the NCI for sharing the Aka/GFP γ-retroviral vector and Dr. David Quach at the Baylor College of Medicine for the DNA template containing ΔEGFR. We also thank the Baylor College of Medicine small animal core and imaging facility for maintaining the animal breeding colonies and for giving access to the IVIS equipment.

## Author contributions

NW conceptualized and designed the study. LL, MN, JE and NW developed the methodology. LL, MN, JE, CO and NW performed the experiments. LL, MN, CO and NW analyzed the data and interpreted the results. LL, MN and NW wrote the original draft of the manuscript. LL, MM and NW secured funding. NW supervised and managed project administration. MN, MM and NW reviewed and edited the manuscript.

## Data Availability

All data and materials used to complete and draw conclusions in this study are presented in the paper, including supplementary information.

---

**Supplementary Figure 1.**
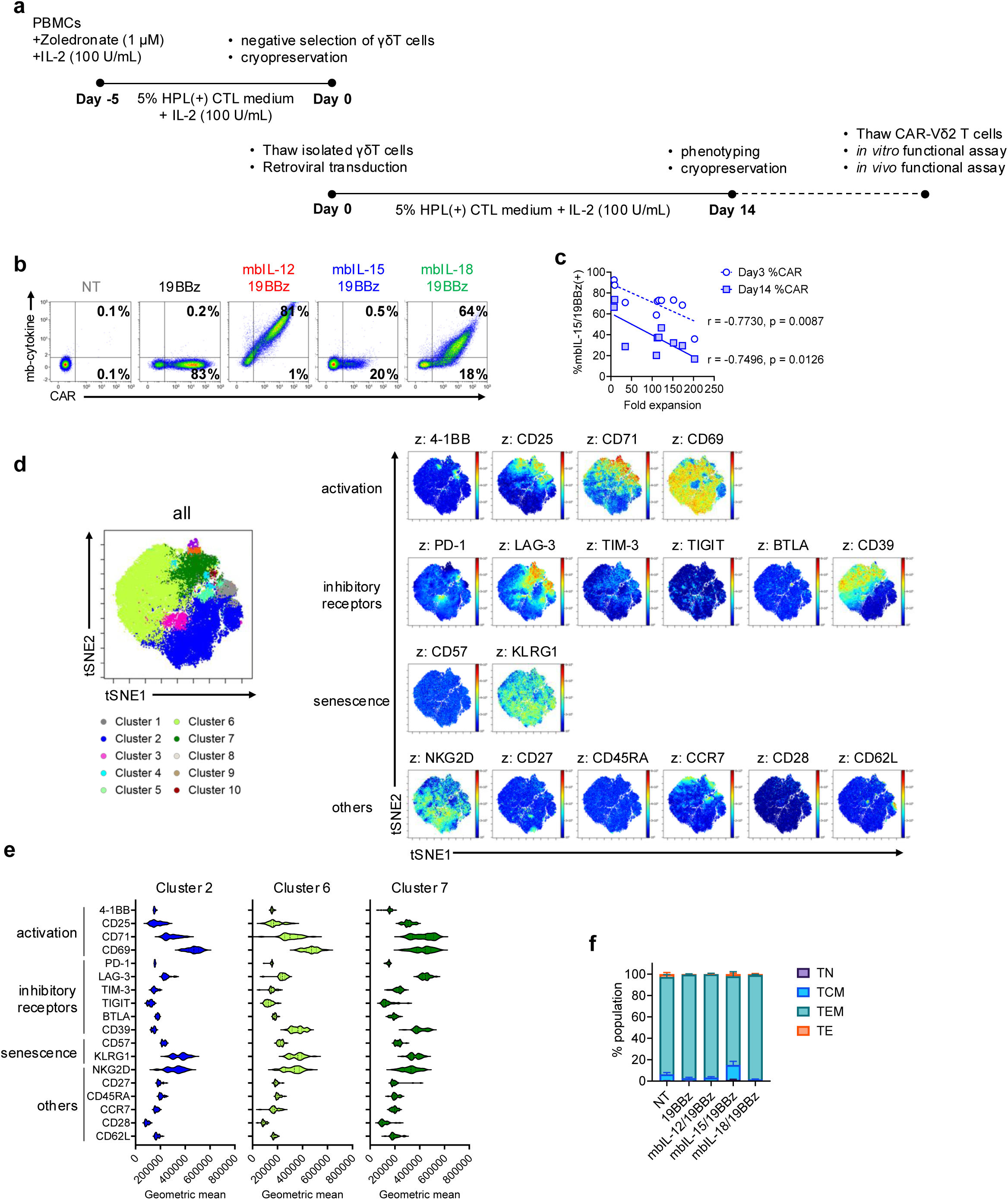
Additional surface phenotype of gene-modified Vδ2 T cells. **a,** Schematic of gene modification and manufacturing of Vδ2 T cells used in this study. **b,** Representative flow plots of surface expression of CAR and mbCyto on day 14 post transduction from 11 independent donors. **c,** Graph showing correlation of % mbIL-15/19BBz(+) cells and total fold cell expansion during 14-day manufacturing. **d,** Visualization of t-distributed stochastic neighbor embedding plots (viSNE) of concatenated events from 5 different cell products with 18 surface phenotype markers followed by FlowSOM metaclustering (cluster 1-10). Left viSNE plot showing all concatenated events including all Vδ2 T cell products with clustering, and right viSNE plots showing overlay of the geometric mean of individual surface marker expression. **e,** Violin plot showing geometric mean of all surface markers expressed in cluster 2, 6 and 7 (mean ± S.E. n=8). **f,** Bar graph summarizing memory ph enotype of each Vδ2 T cell product based on CD27 and CD45RA expression (TN: CD27+CD45RA+, TCM: CD27+CD45RA-, TEM: CD27-CD45RA-, TE: CD27-CD45RA+).

**Supplementary Figure 2.**
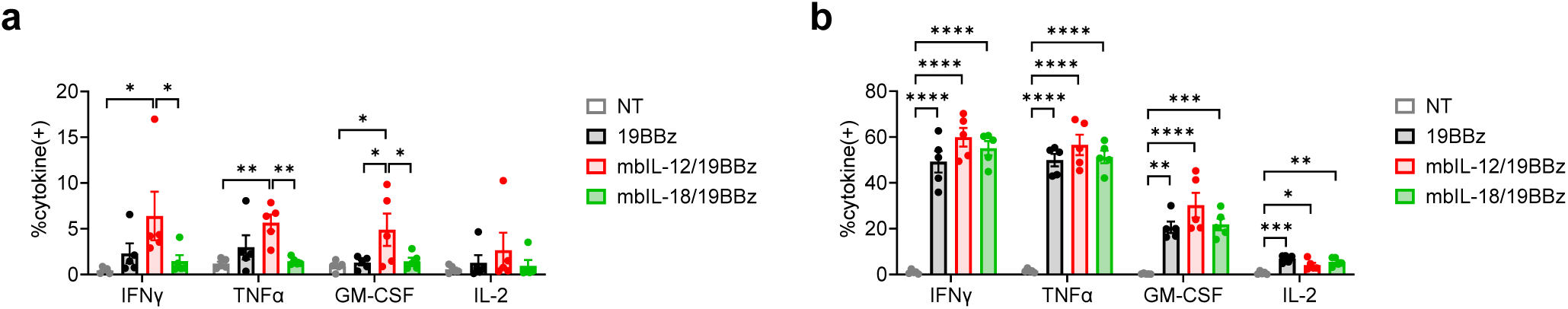
Individual cytokine expression with intracellular cytokine staining. Results of intracellular cytokine staining in each Vδ2 T cell product without antigen stimulation **(a)** or with antigen stimulation **(b)** for 4 hrs (mean ± S.E. n=5). Statistical differences are calculated by One-way ANOVA with Tukey’s multiple comparison in each cytokine (a, b). ^✱^p ≤ 0.05, ^✱✱^p ≤ 0.01, ^✱✱✱^p ≤ 0.001, ^✱✱✱✱^p ≤ 0.0001.

**Supplementary Figure 3.**
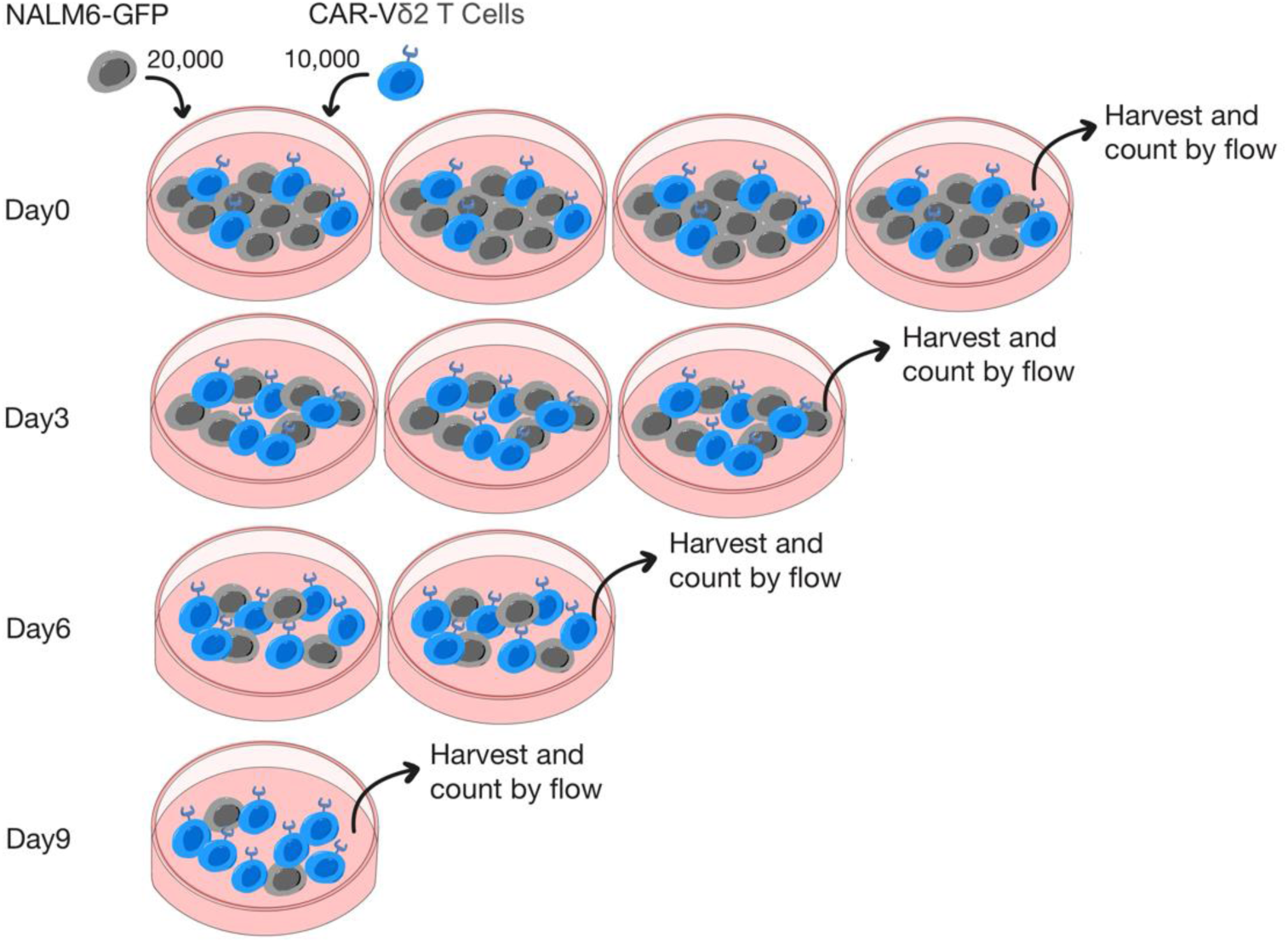
Schematic of *in vitro* 9-day coculture set-up used in this study. Effector cells and target cells are cocultured in indicated effector to target ratio. On the day of set up, total 4 wells are prepared per condition, and every 3 days, cells are harvested from one well and subjected for cell counting on flow cytometry while half of culture medium are replaced for remaining wells for next time point.

**Supplementary Figure 4.**
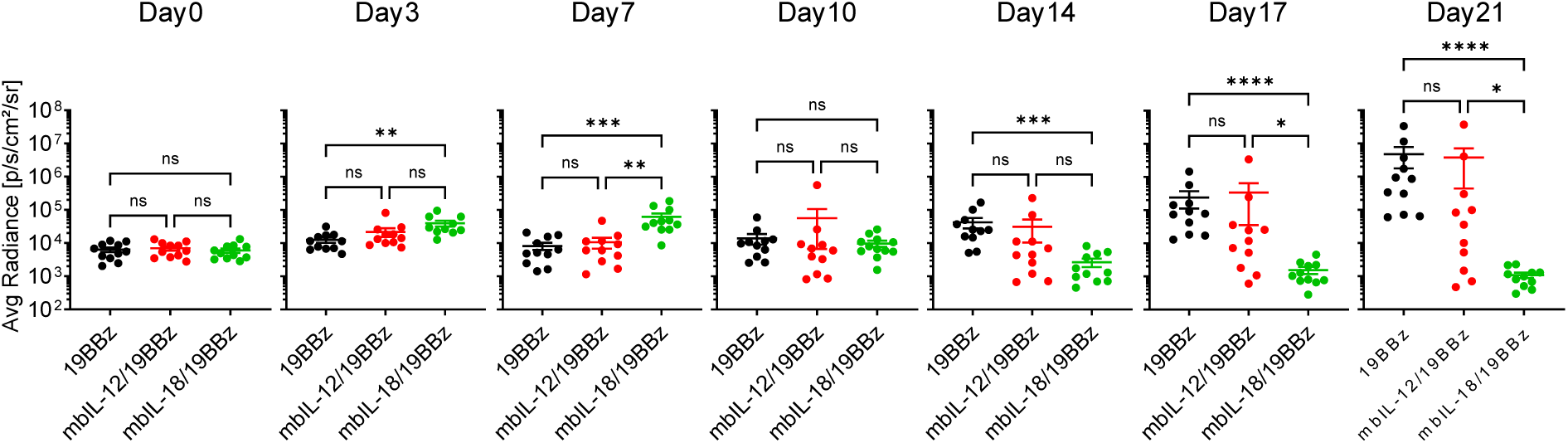
mbIL-18/19BBz Vδ2 T cells delay *in vivo* tumor killing at early time point. Graphs summarizing tumor bioluminescence from individual mice in each treatment group (n=11 in 19BBz and mbIL-12/19BBz, n=12 in mbIL-18/19BBz). Statistical differences are calculated by One-way ANOVA with Tukey’s multiple comparison. ^✱^p ≤ 0.05, ^✱✱^p ≤ 0.01, ^✱✱✱^p ≤ 0.001, ^✱✱✱✱^p ≤ 0.00 01. ns; non-significant.

**Supplementary Figure 5.**
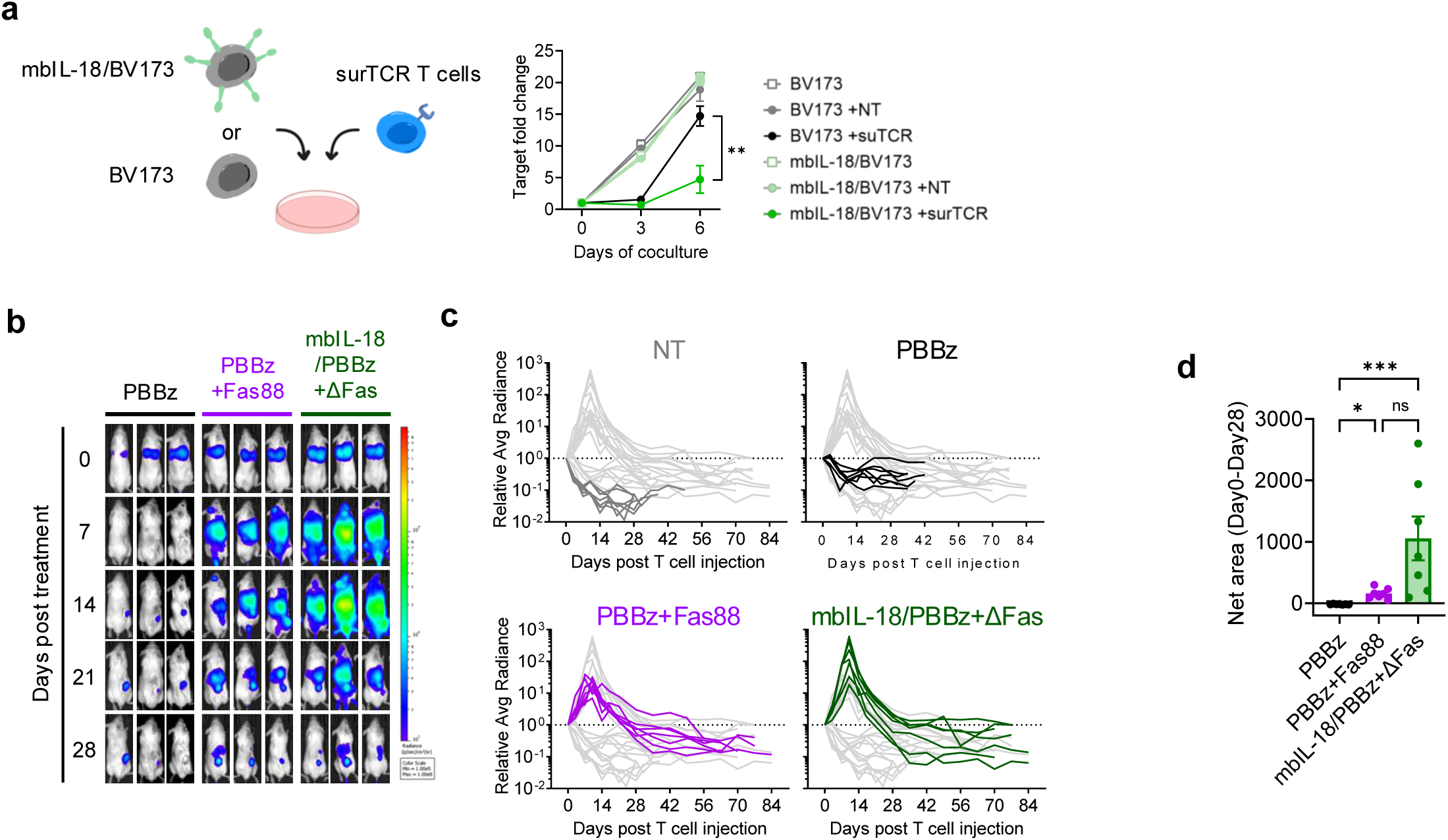
Effect of mbIL-18 trans-presentation on αβT cells and Vδ2 T cell bioluminescence in CFPAC-1 xenograft mouse model. **a,** Schematic of coculture experiments of survivin-specific TCR transduced T cells against BV173 cells with or without mbIL-18 modification (left) and line graph showing target fold change during 6-day coculture (mean ± S.E. n=3). **b,** Representative mice images of Vδ2 T cell bioluminescence at the indicated time points. **c,** Vδ2 T cell expansion and persistence in individual mice over time (n=6 in no treatment and 19BBz, n=5 in 19BBz+Fas88 and mbIL-18/19BBz+ΔFas). **d,** Bar graph showing Net area (baseline = 1) of AkaLuc bioluminescence fold change from day 0 to day 28. Statistical differences are calculated by Paired t test at day 6 (a), One-way ANOVA with Tukey’s multiple comparison (d). ^✱^p ≤ 0.05, ^✱✱^p ≤ 0.01, ^✱✱✱^p ≤ 0.001, ns; non-significant.

**Supplementary Figure 6.**
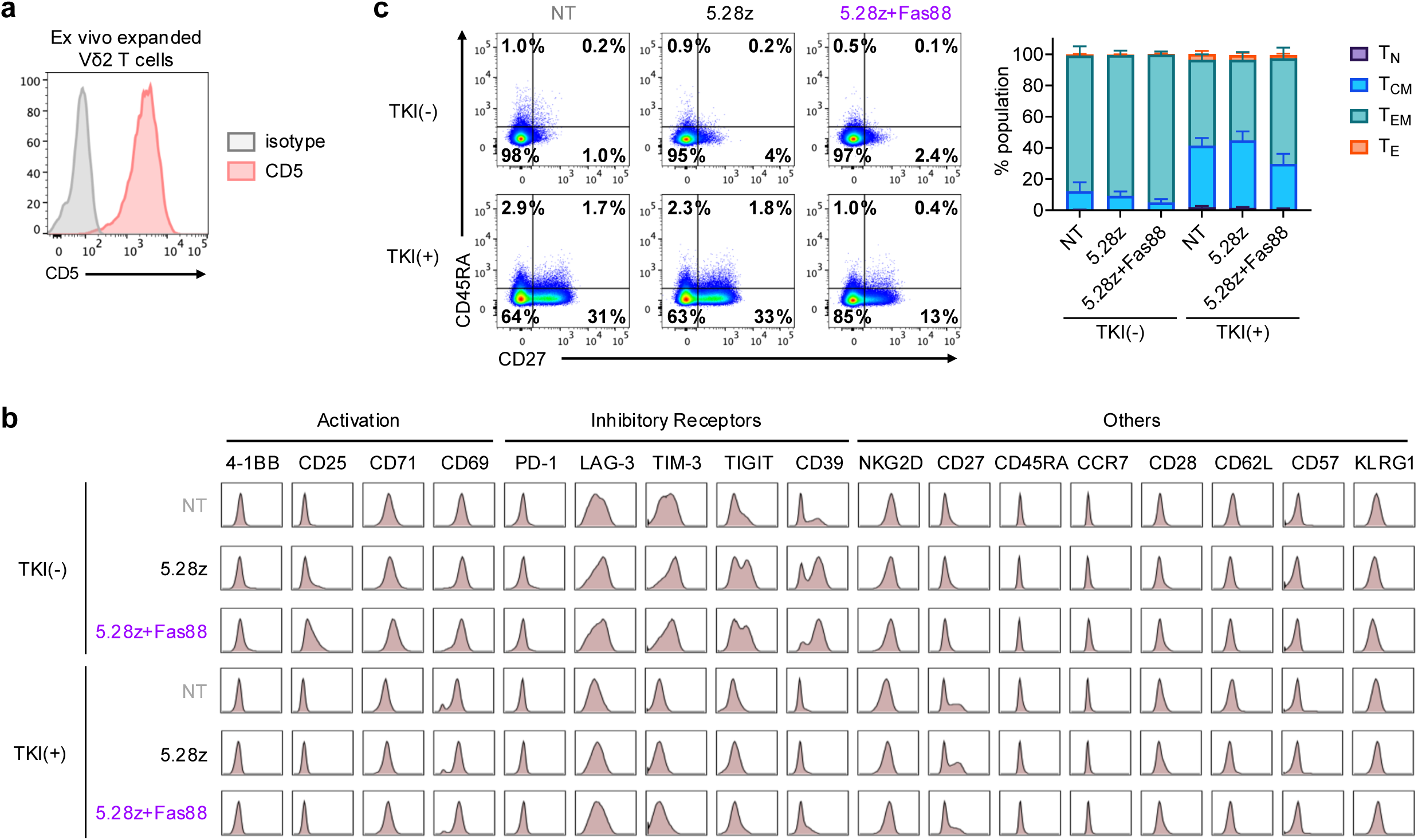
Additional phenotype of CD5.CAR Vδ2 T cells. **a,** Histogram of CD5 surface expression in Vδ2 T cells on the day of retroviral transduction (day 5 of manufacturing process). **b,** Histograms of individual surface marker expression in each of the 6 conditions from visualization of t-distributed stochastic neighbor embedding (viSNE) analysis. **c,** Representative flow plots of the memory phenotype of Vδ2 T cells based on CD27 and CD45RA surface expression (left) and stacked bar graph summarizing memory phenotype in 4 donors/condition (TN: CD27+CD45RA+, TCM: CD27+CD45RA-, TEM: CD27-CD45RA-, TE: CD27-CD45RA+).

**Supplementary Table 1:** List of antibodies used in this study.

| Marker | Fluorochrome | clone | Product # | Vendor |
| --- | --- | --- | --- | --- |
| 2B4 | BV421 | 2-69 | 393504 | BioLegend |
| 4-1BB | BV711 | 4B4-1 | 309832 | BioLegend |
| BTLA | APC/Cy7 | MIH26 | 344518 | BioLegend |
| CCR7 | FITC | 150503 | 561271 | BD Biosciences |
| CD103 | PerCP/Cy5.5 | Ber-ACT8 | 350226 | BioLegend |
| CD16 | BV570 | 3G8 | 302036 | BioLegend |
| CD25 | PE | M-A251 | 555432 | BD Biosciences |
| CD27 | PE/Cy7 | 1A4CD27 | A54823 | Beckman Coulter |
| CD271 (NGFR) | BV421 | C40-1457 | 562562 | BD Biosciences |
| CD271 (NGFR) | BV510 | C40-1457 | 563451 | BD Biosciences |
| CD271 (NGFR) | Pacific Blue | ME20.4 | 345132 | BioLegend |
| CD28 | BV750 | CD28.2 | 302970 | BioLegend |
| CD3 | APC/Fire810 | SK7 | 344858 | BioLegend |
| CD3 | cFluor R685 | SK7 | R7-20345 | CytekBio |
| CD3 | APC/Alexa Fluor750 | UCHT1 | A66329 | Beckman Coulter |
| CD39 | BV480 | TU66 | 746454 | BD Biosciences |
| CD45 | APC/Fire810 | HI30 | 304076 | BioLegend |
| CD45RA | Alexa Fluor 532 | HI100 | 58-0458-42 | Thermo Fischer |
| CD56 | APC Fire 810 | HCD56 | 318356 | BioLegend |
| CD57 | BV605 | QA17A04 | 393304 | BioLegend |
| CD62L | BV785 | GREG-56 | 304830 | BioLegend |
| CD69 | ECD | TP1.55.3 | 6607110 | Beckman Coulter |
| CD71 | PE/Fire 640 | CY1G4 | 334142 | BioLegend |
| Fas | PE | DX2 | 555674 | BD Biosciences |
| Fas ligand | PE/Cy7 | NOK-1 | 306418 | BioLegend |
| FLAG (DYKDDDDK) | BV421 | L5 | 637322 | BioLegend |
| FLAG (DYKDDDDK) | PE/Cy7 | L5 | 637324 | BioLegend |
| G4S Linker | Pacific Blue | E7O2V | 44962S | Cell Signaling Technologies |
| G4S Linker | Alexa Fluor 647 | E7O2V | 69782L | Cell Signaling Technologies |
| GM-CSF | PE/Dazzle 594 | BVD2-21C11 | 502318 | BioLegend |
| IFNγ | V450 | B27 | 560371 | BD Biosciences |
| IL-12 | Biotin |  | BAF219 | R&D System |
| IL-15 | Biotin | BH1543 | 515103 | BioLegend |
| IL-18 | Biotin | 159-12B | D045-6 | MBL Life Science |
| IL-2 | APC/Fire750 | MQ1-17H12 | 500352 | BioLegend |
| KLRG1 | PerCP eFluor 710 | 13F12F2 | 46-9488-42 | Thermo Fischer |
| LAG-3 | PE-Cy5 | 11C3C65 | 369346 | BioLegend |
| NKG2D | APC | 1D11 | 320808 | BioLegend |
| PD-1 | BV650 | EH12.2H7 | 329950 | BioLegend |
| Phospho-p38 MAPK (pT180/pY182) | Alexa Fluor 647 | 36/p38 | 612595 | BD Biosciences |
| Phospho-STAT4 (pY693) | Alexa Fluor 647 | 38/p-Stat4 | 558137 | BD Biosciences |
| Phospho-STAT5 (Tyr694) | PE-Cy7 | A17016B.Rec | 936908 | BioLegend |
| Streptavidin | APC |  | 554067 | BD Bioscience |
| TIGIT | PE-Fire 810 | A15153G | 372745 | BioLegend |
| TIM-3 | Alexa Fluor 700 | F38-2E2 | 56-3109-42 | Thermo Fischer |
| TNFα | PE-Cy7 | MAb11 | 557647 | BD Biosciences |
| Vδ1TCR | VioBlue | REA173 \| R9.12 | 130-120-443 | Milteny Biotec |
| Vδ1TCR | Biotin | REA173 \| R9.12 | 130-120-439 | Milteny Biotec |
| Vδ2TCR | BV510 | B6 | 331432 | BioLegend |
| Vδ2TCR | PE | B6 | 555739 | BD Biosciences |
| Vδ2TCR | Biotin | REA771 \| 123R3 | 130-111-008 | Milteny Biotec |
| $\alpha\beta$TCR | Biotin | REA652 \| BW242/412 | 130-113-537 | Milteny Biotec |
| $\alpha\beta$TCR | FITC | WT31 | 347773 | BD Biosciences |
| Phospho-NF-κB p65 (Ser536) | | 93H1 | #3033 | Cell Signaling Technologies |
| Phospho-IKKα/$\beta$ (Ser176/180) | | 16A6 | #2697 | Cell Signaling Technologies |
| Phospho-p38 MAPK (Thr180/Tyr182) |  |  | #9211 | Cell Signaling Technologies |
| GAPDH |  | 6C5 | sc-32233 | Santa Cruz |
| Goat anti-Rabbit IgG Secondary Antibody | IRDye® 800CW |  | 926-32211 | LICORBio |
| Goat Anti-Mouse IgG Secondary Antibody | IRDye® 680RD |  | 926-68070 | LICORBio |

## References

1 Hu, Y. et al. γδ T cells: origin and fate, subsets, diseases and immunotherapy. Signal Transduct Target Ther 8, 434, doi:10.1038/s41392-023-01653-8 (2023).

2 Gu, S., Borowska, M. T., Boughter, C. T. & Adams, E. J. Butyrophilin3A proteins and Vγ9Vδ2 T cell activation. Semin Cell Dev Biol 84, 65–74, doi:10.1016/j.semcdb.2018.02.007 (2018).

3 Rigau, M. et al. Butyrophilin 2A1 is essential for phosphoantigen reactivity by γδ T cells. Science 367, doi:10.1126/science.aay5516 (2020).

4 Liu, Y. & Zhang, C. The Role of Human γδ T Cells in Anti-Tumor Immunity and Their Potential for Cancer Immunotherapy. Cells 9, doi:10.3390/cells9051206 (2020).

5 Rozenbaum, M. et al. Gamma-Delta CAR-T Cells Show CAR-Directed and Independent Activity Against Leukemia. Front Immunol 11, 1347, doi:10.3389/fimmu.2020.01347 (2020).

6 Thomas, P., Paris, P. & Pecqueur, C. Arming Vδ2 T Cells with Chimeric Antigen Receptors to Combat Cancer. Clin Cancer Res 30, 3105–3116, doi:10.1158/1078-0432.CCR-23-3495 (2024).

7 Lee, D. et al. Unlocking the potential of allogeneic Vδ2 T cells for ovarian cancer therapy through CD16 biomarker selection and CAR/IL-15 engineering. Nat Commun 14, 6942, doi:10.1038/s41467-023-42619-2 (2023).

8 Ma, L., Feng, Y. & Zhou, Z. A close look at current γδ T-cell immunotherapy. Front Immunol 14, 1140623, doi:10.3389/fimmu.2023.1140623 (2023).

9 Barisa, M., Nattress, C., Fowler, D., Anderson, J. & Fisher, J. in γδT Cell Cancer Immunotherapy (ed Marta Barisa) 103–153 (Academic Press, 2025).

10 Revesz, I. A., Joyce, P., Ebert, L. M. & Prestidge, C. A. Effective γδ T-cell clinical therapies: current limitations and future perspectives for cancer immunotherapy. Clin Transl Immunology 13, e1492, doi:10.1002/cti2.1492 (2024).

11 Gan, Y. H., Lui, S. S. & Malkovsky, M. Differential susceptibility of naïve and activated human gammadelta T cells to activation-induced cell death by T-cell receptor cross-linking. Mol Med 7, 636–643 (2001).

12 Frieling, J. S. et al. γδ-Enriched CAR-T cell therapy for bone metastatic castrate-resistant prostate cancer. Sci Adv 9, eadf0108, doi:10.1126/sciadv.adf0108 (2023).

13 Wang, Y. et al. CAR-Modified Vγ9Vδ2 T Cells Propagated Using a Novel Bisphosphonate Prodrug for Allogeneic Adoptive Immunotherapy. Int J Mol Sci 24, doi:10.3390/ijms241310873 (2023).

14 Deniger, D. C. et al. Bispecific T-cells expressing polyclonal repertoire of endogenous γδ T-cell receptors and introduced CD19-specific chimeric antigen receptor. Mol Ther 21, 638–647, doi:10.1038/mt.2012.267 (2013).

15 Zhai, X. et al. MUC1-Tn-targeting chimeric antigen receptor-modified Vγ9Vδ2 T cells with enhanced antigen-specific anti-tumor activity. Am J Cancer Res 11, 79–91 (2021).

16 Song, Y., Liu, Y., Teo, H. Y. & Liu, H. Targeting Cytokine Signals to Enhance γδT Cell-Based Cancer Immunotherapy. Front Immunol 13, 914839, doi:10.3389/fimmu.2022.914839 (2022).

17 Jia, Z. et al. IL12 immune therapy clinical trial review: Novel strategies for avoiding CRS-associated cytokines. Front Immunol 13, 952231, doi:10.3389/fimmu.2022.952231 (2022).

18 Mo, F. et al. Human platelet lysate enhances in vivo activity of CAR-Vδ2 T cells by reducing cellular senescence and apoptosis. Cytotherapy 26, 858–868, doi:10.1016/j.jcyt.2024.03.006 (2024).

19 Muntasell, A., Magri, G., Pende, D., Angulo, A. & López-Botet, M. Inhibition of NKG2D expression in NK cells by cytokines secreted in response to human cytomegalovirus infection. Blood 115, 5170–5179, doi:10.1182/blood-2009-11-256479 (2010).

20 Arber, C. et al. Survivin-specific T cell receptor targets tumor but not T cells. J Clin Invest 125, 157–168, doi:10.1172/JCI75876 (2015).

21 Ma, R. et al. Chimeric antigen receptor-induced antigen loss protects CD5.CART cells from fratricide without compromising on-target cytotoxicity. Cell Rep Med 5, 101628, doi:10.1016/j.xcrm.2024.101628 (2024).

22 Hill, L. C. et al. Antitumor efficacy and safety of unedited autologous CD5.CAR T cells in relapsed/refractory mature T-cell lymphomas. Blood 143, 1231–1241, doi:10.1182/blood.2023022204 (2024).

23 Watanabe, N. et al. Feasibility and preclinical efficacy of CD7-unedited CD7 CAR T cells for T cell malignancies. Mol Ther 31, 24–34, doi:10.1016/j.ymthe.2022.09.003 (2023).

24 Yuan, M. et al. Advancements in γδT cell engineering: paving the way for enhanced cancer immunotherapy. Front Immunol 15, 1360237, doi:10.3389/fimmu.2024.1360237 (2024).

25 Xu, Y. et al. Allogeneic Vγ9Vδ2 T-cell immunotherapy exhibits promising clinical safety and prolongs the survival of patients with late-stage lung or liver cancer. Cell Mol Immunol 18, 427–439, doi:10.1038/s41423-020-0515-7 (2021).

26 Teo, H. Y. et al. IL12/18/21 Preactivation Enhances the Antitumor Efficacy of Expanded γδT Cells and Overcomes Resistance to Anti-PD-L1 Treatment. Cancer Immunol Res 11, 978–999, doi:10.1158/2326-6066.CIR-21-0952 (2023).

27 Schilbach, K. et al. In the Absence of a TCR Signal IL-2/IL-12/18-Stimulated γδ T Cells Demonstrate Potent Anti-Tumoral Function Through Direct Killing and Senescence Induction in Cancer Cells. Cancers (Basel) 12, doi:10.3390/cancers12010130 (2020).

28 Caccamo, N. et al. Differentiation, phenotype, and function of interleukin-17-producing human Vγ9Vδ2 T cells. Blood 118, 129–138, doi:10.1182/blood-2011-01-331298 (2011).

29 Barjon, C. et al. IL-21 promotes the development of a CD73-positive Vγ9Vδ2 T cell regulatory population. Oncoimmunology 7, e1379642, doi:10.1080/2162402X.2017.1379642 (2017).

30 Lafont, V. et al. Plasticity of γδ T Cells: Impact on the Anti-Tumor Response. Front Immunol 5, 622, doi:10.3389/fimmu.2014.00622 (2014).

31 Wesch, D., Glatzel, A. & Kabelitz, D. Differentiation of resting human peripheral blood gamma delta T cells toward Th1- or Th2-phenotype. Cell Immunol 212, 110–117, doi:10.1006/cimm.2001.1850 (2001).

32 Li, W. et al. Effect of IL-18 on expansion of gammadelta T cells stimulated by zoledronate and IL-2. J Immunother 33, 287–296, doi:10.1097/CJI.0b013e3181c80ffa (2010).

33 Koneru, M., Purdon, T. J., Spriggs, D., Koneru, S. & Brentjens, R. J. IL-12 secreting tumor-targeted chimeric antigen receptor T cells eradicate ovarian tumors. Oncoimmunology 4, e994446, doi:10.4161/2162402X.2014.994446 (2015).

34 Liu, Y. et al. Armored Inducible Expression of IL-12 Enhances Antitumor Activity of Glypican-3-Targeted Chimeric Antigen Receptor-Engineered T Cells in Hepatocellular Carcinoma. J Immunol 203, 198–207, doi:10.4049/jimmunol.1800033 (2019).

35 Lee, E. H. J. et al. Antigen-dependent IL-12 signaling in CAR T cells promotes regional to systemic disease targeting. Nat Commun 14, 4737, doi:10.1038/s41467-023-40115-1 (2023).

36 Fowler, D. et al. Payload-delivering engineered γδ T cells display enhanced cytotoxicity, persistence, and efficacy in preclinical models of osteosarcoma. Sci Transl Med 16, eadg9814, doi:10.1126/scitranslmed.adg9814 (2024).

37 Canestrari, E., Steidinger, H. R., McSwain, B., Charlebois, S. J. & Dann, C. T. Human Platelet Lysate Media Supplement Supports Lentiviral Transduction and Expansion of Human T Lymphocytes While Maintaining Memory Phenotype. J Immunol Res 2019, 3616120, doi:10.1155/2019/3616120 (2019).

38 Peters, C. et al. TGF-β enhances the cytotoxic activity of Vδ2 T cells. Oncoimmunology 8, e1522471, doi:10.1080/2162402X.2018.1522471 (2019).

39 Silva, J. A. et al. Central memory-enriched Vγ9Vδ2 γδ T cells via TGF-β expansion demonstrate enhanced. Front Immunol 16, 1657760, doi:10.3389/fimmu.2025.1657760 (2025).

40 Casetti, R. et al. Cutting edge: TGF-beta1 and IL-15 Induce FOXP3+ gammadelta regulatory T cells in the presence of antigen stimulation. J Immunol 183, 3574–3577, doi:10.4049/jimmunol.0901334 (2009).

41 Jaspers, J. E. et al. IL-18-secreting CAR T cells targeting DLL3 are highly effective in small cell lung cancer models. J Clin Invest 133, doi:10.1172/JCI166028 (2023).

42 Li, Y. R., Zhu, Y. & Yang, L. IL-18 revives dysfunctional CAR-T cells. Trends Cancer 11, 923–926, doi:10.1016/j.trecan.2025.06.008 (2025).

43 Hu, B. et al. Augmentation of Antitumor Immunity by Human and Mouse CAR T Cells Secreting IL-18. Cell Rep 20, 3025–3033, doi:10.1016/j.celrep.2017.09.002 (2017).

44 Avanzi, M. P. et al. Engineered Tumor-Targeted T Cells Mediate Enhanced Anti-Tumor Efficacy Both Directly and through Activation of the Endogenous Immune System. Cell Rep 23, 2130–2141, doi:10.1016/j.celrep.2018.04.051 (2018).

45 Svoboda, J. et al. Enhanced CAR T-Cell Therapy for Lymphoma after Previous Failure. N Engl J Med 392, 1824–1835, doi:10.1056/NEJMoa2408771 (2025).

46 Geyer, M. B. et al. CD371-Targeted CAR T cells Secreting Interleukin-18 Exhibit Robust Expansion and Clear Refractory Acute Myeloid Leukemia. Blood, doi:10.1182/blood.2025029532 (2025).

47 Hull, C. M. et al. Granzyme B-activated IL18 potentiates αβ and γδ CAR T cell immunotherapy in a tumor-dependent manner. Mol Ther 32, 2373–2392, doi:10.1016/j.ymthe.2024.05.013 (2024).

48 Blokon-Kogan, D. et al. Membrane anchored IL-18 linked to constitutively active TLR4 and CD40 improves human T cell antitumor capacities for adoptive cell therapy. Journal for ImmunoTherapy of Cancer 10, e001544, doi:10.1136/jitc-2020-001544 (2022).

49 Yamamoto, T. N. et al. T cells genetically engineered to overcome death signaling enhance adoptive cancer immunotherapy. J Clin Invest 129, 1551–1565, doi:10.1172/JCI121491 (2019).

50 Yi, F. et al. CAR-engineered lymphocyte persistence is governed by a FAS ligand-FAS autoregulatory circuit. Nat Cancer 6, 1638–1655, doi:10.1038/s43018-025-01009-x (2025).

51 Anderson, K. G. et al. Engineering adoptive T cell therapy to co-opt Fas ligand-mediated death signaling in ovarian cancer enhances therapeutic efficacy. J Immunother Cancer 10, doi:10.1136/jitc-2021-003959 (2022).

52 McKenzie, C. et al. Novel Fas-TNFR chimeras that prevent Fas ligand-mediated kill and signal synergistically to enhance CAR T cell efficacy. Mol Ther Nucleic Acids 32, 603–621, doi:10.1016/j.omtn.2023.04.017 (2023).

53 Oda, S. K. et al. A Fas-4-1BB fusion protein converts a death to a pro-survival signal and enhances T cell therapy. J Exp Med 217, doi:10.1084/jem.20191166 (2020).

54 Sahillioglu, A. C. & Schumacher, T. N. Safety switches for adoptive cell therapy. Curr Opin Immunol 74, 190–198, doi:10.1016/j.coi.2021.07.002 (2022).

55 Jan, M. et al. Reversible ON- and OFF-switch chimeric antigen receptors controlled by lenalidomide. Sci Transl Med 13, doi:10.1126/scitranslmed.abb6295 (2021).

56 Straathof, K. C. et al. An inducible caspase 9 safety switch for T-cell therapy. Blood 105, 4247–4254, doi:10.1182/blood-2004-11-4564 (2005).

57 Mamonkin, M., Rouce, R. H., Tashiro, H. & Brenner, M. K. A T-cell-directed chimeric antigen receptor for the selective treatment of T-cell malignancies. Blood 126, 983–992, doi:10.1182/blood-2015-02-629527 (2015).

58 Watanabe, N. et al. Fine-tuning the CAR spacer improves T-cell potency. Oncoimmunology 5, e1253656, doi:10.1080/2162402X.2016.1253656 (2016).

59 Marks-Konczalik, J. et al. IL-2-induced activation-induced cell death is inhibited in IL-15 transgenic mice. Proc Natl Acad Sci U S A 97, 11445–11450, doi:10.1073/pnas.200363097 (2000).

60 Zhang, L. et al. Enhanced efficacy and limited systemic cytokine exposure with membrane-anchored interleukin-12 T-cell therapy in murine tumor models. J Immunother Cancer 8, doi:10.1136/jitc-2019-000210 (2020).

61 Pan, W. Y. et al. Cancer immunotherapy using a membrane-bound interleukin-12 with B7-1 transmembrane and cytoplasmic domains. Mol Ther 20, 927–937, doi:10.1038/mt.2012.10 (2012).

62 Jin, B. Y. et al. Engineered T cells targeting E7 mediate regression of human papillomavirus cancers in a murine model. JCI Insight 3, doi:10.1172/jci.insight.99488 (2018).

63 Narayanan, P. et al. A composite MyD88/CD40 switch synergistically activates mouse and human dendritic cells for enhanced antitumor efficacy. J Clin Invest 121, 1524–1534, doi:10.1172/JCI44327 (2011).

64 Torres Chavez, A. G., et al. A dual-luciferase bioluminescence system for the assessment of cellular therapies. Mol Ther Oncol 32, 200763, doi:10.1016/j.omton.2024.200763 (2024).

65 Gomes-Silva, D. et al. Tonic 4-1BB Costimulation in Chimeric Antigen Receptors Impedes T Cell Survival and Is Vector-Dependent. Cell Rep 21, 17–26, doi:10.1016/j.celrep.2017.09.015 (2017).

66 Watanabe, N. & McKenna, M. K. Generation of CAR T-cells using γ-retroviral vector. Methods Cell Biol 167, 171–183, doi:10.1016/bs.mcb.2021.06.014 (2022).

67 Babicki, S. et al. Heatmapper: web-enabled heat mapping for all. Nucleic Acids Res 44, W147–153, doi:10.1093/nar/gkw419 (2016).

